# Hyperexcitability in human *MECP2* null neuronal networks manifests as calcium-dependent reverberating super bursts

**DOI:** 10.1101/2023.09.12.557388

**Authors:** Kartik S Pradeepan, Fraser P McCready, Wei Wei, Milad Khaki, Wenbo Zhang, Michael W Salter, James Ellis, Julio Martinez-Trujillo

**Affiliations:** Graduate Program in Neuroscience, Schulich School of Medicine and Dentistry, Western University, London, ON, Canada; Robarts Research Institute, Western University, London, ON, Canada; Department of Molecular Genetics, University of Toronto, Toronto, ON, Canada; Developmental & Stem Cell Biology, The Hospital for Sick Children, Toronto, ON, Canada; Neuroscience & Mental Health, The Hospital for Sick Children, Toronto, ON, Canada; Department of Physiology, University of Toronto, ON, Canada; Brain and Mind Institute, Western University, London, ON, Canada; Department of Physiology and Pharmacology, Schulich School of Medicine and Dentistry, Western University, London, ON, Canada; Department of Psychiatry, Schulich School of Medicine and Dentistry, Western University, London, ON, Canada

**Keywords:** network reverberations, reverberating super bursts, super bursts, multielectrode array, burst detection, Rett syndrome, human stem cell derived neurons, electrophysiology, neurodevelopmental disorder

## Abstract

Rett syndrome (RTT) patients show abnormal developmental trajectories including loss of language and repetitive hand movements but also have signs of cortical hyperexcitability such as seizures. RTT is predominantly caused by mutations in *MECP2* and can be modelled in vitro using human stem cell-derived neurons. *MECP2* null excitatory neurons are smaller in soma size and have reduced synaptic connectivity but are also hyperexcitable, due to higher input resistance, which increases the chance to evoke action potentials with a given depolarized current. Few studies examine how single neuron activity integrates into neuronal networks during human development. Paradoxically, networks of *MECP2* null neurons show a decrease in the frequency of bursting patterns consistent with synaptic hypoconnectivity, but no hyperexcitable network events have been reported. Here, we show that *MECP2* null neurons have an increase in the frequency of a network event described as reverberating super bursts (RSBs) relative to isogenic controls. RSBs can be mistakenly called as a single long duration burst by standard burst detection algorithms. However, close examination revealed an initial large amplitude network burst followed by high frequency repetitive low amplitude mini-bursts. Using a custom burst detection algorithm, we unfolded the multi-burst structure of RSBs revealing that *MECP2* null networks increased the total number of bursts relative to isogenic controls. Application of the Ca^2+^ chelator EGTA-AM selectively eliminated RSBs and rescued the network burst phenotype relative to the isogenic controls. Our results indicate that during early development, *MECP2* null neurons are hyperexcitable and produce hyperexcitable networks. This may predispose them to the emergence of hyper-synchronic states that potentially translate into seizures. Network hyperexcitability is dependent on asynchronous neurotransmitter release driven by pre-synaptic Ca^2+^ and can be rescued by EGTA-AM to restore typical network dynamics.

**HIGHLIGHTS:**

1. Reverberating super-bursts (RSBs) follow a stereotypic form of a large initial network burst followed by several smaller amplitude high-frequency mini-bursts.
2. RSBs occur more often in *MECP2* null excitatory networks.
3. *MECP2* null excitatory networks with increased RSBs show a hyperexcitable network burst phenotype relative to isogenic controls.
4. The calcium chelator, EGTA-AM, decreases RSBs and rescues the dynamics of *MECP2* null hyperexcitable networks.

## INTRODUCTION

Mutations of the X-linked gene methyl-CpG-binding protein 2 (*MECP2*), coding for a global transcriptional regulator that is critically involved in the development of the nervous system, cause Rett syndrome (RTT)^1–4^. RTT patients show normal early development until 6-18 months of age when acquired language is lost and repetitive hand movements appear, with epilepsy in up to 60-80% of cases epilepsy^2,4–6^. The loss of MECP2 generates an opaque and complex cascade of structural and functional changes in neurons during development. How such changes produce features of the RTT clinical phenotype such as neuronal network hyperexcitability and seizures remains unclear.

An emerging tool to study early human neuronal network development is in vitro multielectrode array (MEA) electrophysiological recordings^7–10^. Neuronal cultures or organoids from patients or controls are plated overtop a grid of electrodes that measure the activity of neurons in the vicinity of each electrode^8^. Stereotypical patterns of spontaneous action potential firing first emerge in single neurons^7^. Later, as neurons wire together, their activity progresses to patterns of correlated firing known as network bursts^11–15^, which are thought to shape the development of synapses^16^. Activity patterns in mutant networks, such as those derived from RTT patients that deviate from those observed in control networks, may reflect altered normal development and progression to neurological disorders^17^. Here we use MEA recordings to investigate activity patterns during the early stages of development in *MECP2* mutant networks and isogenic controls. The process of spontaneous activity in single neurons developing into more complex correlated network activity is driven by a combination of both intrinsic (e.g., neuronal maturation – as neurons mature, their membrane properties evolve and their capacity to generate action potentials and therefore functional connections with other neurons increase) and extrinsic factors (e.g., activity-dependent plasticity in synapses, and neuromodulators)^26,27^. One prevalent activity pattern in stem cell-derived neuronal networks is bursting. Bursts are a type of complex spontaneous activity, referring to a group of action potentials fired in rapid succession above a low baseline firing rate followed by a period of return to baseline^18^. During development, single neuron bursts synchronize forming network bursts – indicative of neurons connecting within a network ^18–21^. As a result, network burst features, such as network burst duration and frequency, are commonly reported phenotypic metrics^8^.

In this current study, we closely examined the bursting activity of stem cell-derived *MECP2* mutant neuronal networks. Animal models have suggested that *MECP2* mutations disrupt synaptic function^1,22,23^. Supporting this hypothesis, rodent models show delayed maturation of functional synaptic connections^4,24^. Additionally, we previously reported that human stem cell-derived excitatory neurons harbouring an *MECP2* null mutation produce morphologically and functionally hypo-connected neurons with reduced spontaneous action potential numbers^7^. The same MECP2 null neurons, when isolated from the network, showed changes in intrinsic properties such as increased input resistance which is generally inversely related to rheobase, suggesting hyperexcitable neurons^7^. Together with decreased capacitance^7^, input resistance jointly contributes to the neuron’s time course and response to current injections. Paradoxically, networks of *MECP2* null neurons show a decrease in the frequency of network bursts^7^. These findings are also discordant with the increase in hyperexcitability and hyper-synchronicity that leads to seizures in many RTT patients^5^.

In the present study, we aim to clarify these issues by recording the activity of human stem cell-derived single neurons and networks carrying *MECP2* mutations. We found that *MECP2* mutant networks exhibit a temporally complex bursting pattern termed reverberating super-burst (RSB), which consists of an initial large amplitude burst followed by lower amplitude but high-frequency mini-bursts. The latter are undetected using standard burst detection algorithms. RSBs lead to an increase in the frequency of single network bursts in *MECP2* null networks relative to isogenic controls. We finally explored mechanisms of RSB generation. Previous literature has implicated inhibitory neurotransmission^25,26^ and calcium dynamics^21,25,27–29^ in modulating reverberation-like activity. As such, we applied bicuculline (a GABA_A_ receptor antagonist) and EGTA-AM (a slow-kinetic, membrane-permeable Ca^2+^ chelator) to reveal what dynamics may be at play in our excitatory networks. We demonstrated that EGTA-AM decreases RSB frequency, thereby rescuing network bursting dynamics in *MECP2* null networks.

## RESULTS

### A hyperexcitability phenotype in MECP2 null single neurons

Electrophysiological data recorded by Mok et al. (2022) from *MECP2* mutant excitatory neurons and neuronal networks including their isogenic controls were obtained from NDD-Ephys-dB (see Materials and Methods and ^7^). This newly created database facilitates sharing of raw data from electrophysiology experiments and incorporates a toolbox for improved exploration and visualization of results. In brief, Mok et al. (2022) generated excitatory neurons from three isogenic pairs of female pluripotent stem cell lines (WIBR3, PGPC14, and CLT) using the *Ngn2* protocol (fig. 1, b and ^7^). The *MECP2* null pairs were derived by gene editing of the WIBR3 human embryonic stem cell (ESC) line or the PGPC14 healthy induced pluripotent stem cell (iPSC) line. CLT iPSC lines were derived from an atypical RTT patient with the milder phenotype of the preserved speech variant (heterozygous L124W missense mutation) and express only the mutant or the wildtype allele due to X-chromosome inactivation. Stem cell-derived *MECP2* null excitatory neurons from WIBR3 and PGPC14 have smaller cell bodies and dendritic arbours relative to isogenic controls, but CLT neurons have very minor changes in dendritic branching only^7^.

**Figure 1.**
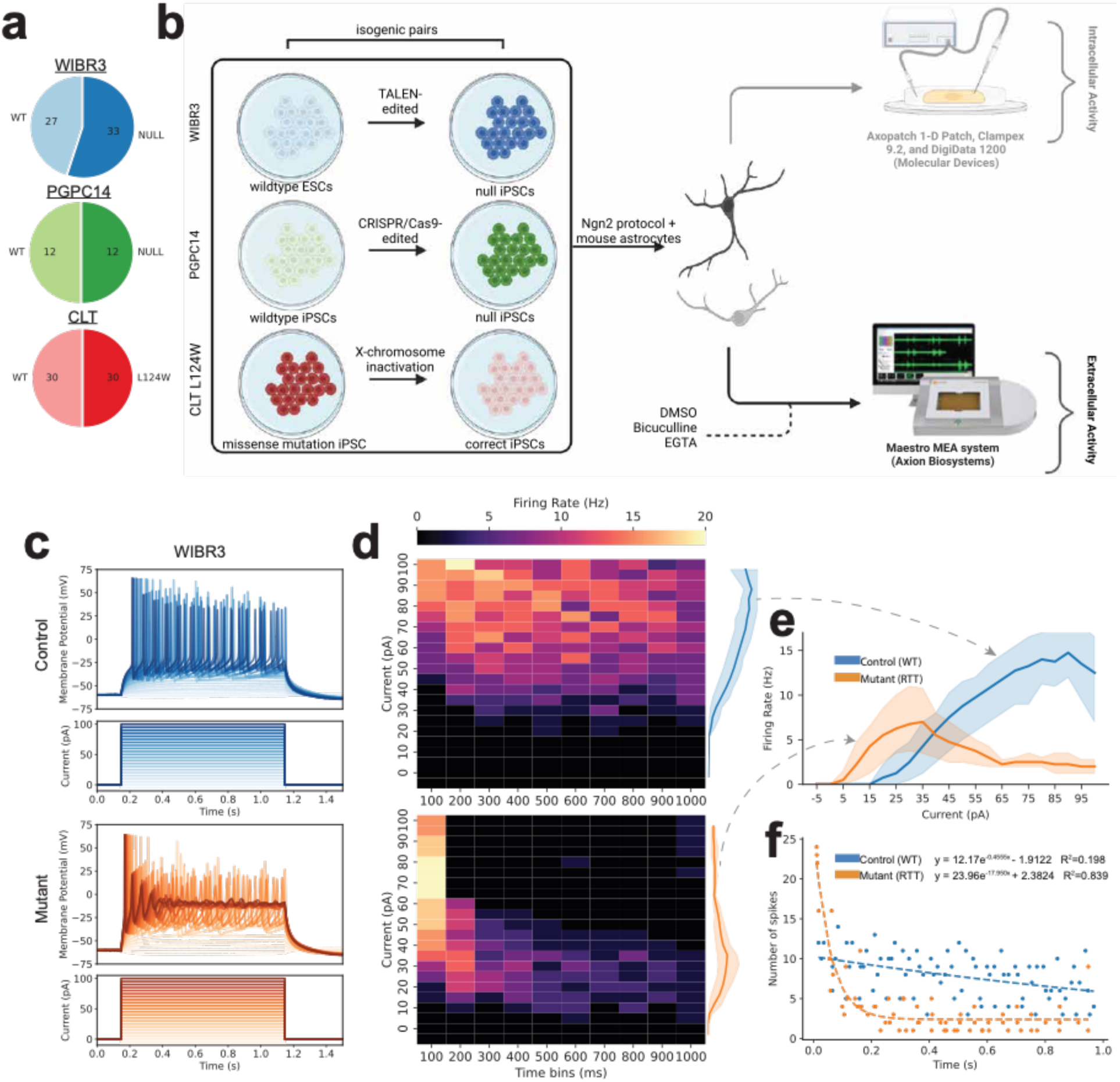
Study design and hyperexcitability in *MECP2* null single neurons. (a) Proportion of wildtype and *MECP2* mutant human stem cell lines used from Mok et al., (2022) which can be found on NDD-Ephys-dB. (b) Schematic overview. Three female cell lines (WIBR3-unaffected embryonic stem cell; PGPC14-unaffected induced pluripotent stem cell (iPSC); CLT L124W-affected patient-derived iPSC) were used to generate 3 isogenic pairs for a total of 6 lines. Each line was differentiated into excitatory neurons using a rapid single-step *Ngn2* differentiation protocol and plated overtop mouse astrocytes. Intracellular single-neuron activity was measured between weeks 4-5. Extracellular population activity was measured for 5 minutes weekly for 6 weeks. At week 5, pharmacological treatment with bicuculline and EGTA-AM was performed on WIBR3. (c) Representative 1-second square pulse current clamp traces with 5pA increments. (d) Firing rate heatmap of WIBR3 neurons with 100ms time bins. (e) Input response curve of average WIBR3 intracellular activity. WIBR3 null single neurons exhibit lower rheobase indicating hyper-excitability. (f) Exponential decay curve fit of the number of spikes per cell across sweeps over 1-second square pulse. WIBR3 null single neurons exhibit greater adaptation indicated by steeper decay.

Our re-analysis of the data confirms that WIBR3 *MECP2* null neurons show a lower rheobase than isogenic controls, i.e., they fire action potentials at lower stimulation currents than isogenic controls (fig. 1c-e). They also show a progressive increase in firing rate that peaks and then declines with the enhanced intensity of the stimulus (fig. 1d-e). On the other hand, isogenic controls show higher rheobase but a sustained increase in firing frequency to much higher than the *MECP2* null and then plateau in response to the enhanced intensity of the stimulus (fig. 1d, e, and see ^7^). Spike rate adaptation, quantified as the absolute value of the slope of an exponential decay function fitted to the firing rate as a function of stimulation current intensity, is stronger in *MECP2* null neurons compared to controls (fig. 1f). These findings indicate that *MECP2* null single neurons, although more hyperexcitable than controls, reach lower peak firing rates and undergo stronger spike rate adaptation relative to isogenic controls.

### Bursting phenotypes in MECP2 mutant neuronal networks

One seemingly contradictory finding from the earlier study that used MEA recordings was a decrease in network burst frequency and increased network burst duration in *MECP2* null networks relative to isogenic control networks^7^. This was puzzling because the WIBR3 *MECP2* null single neurons were more excitable and showed faster adaptation than the controls, yet the MEA data revealed networks with longer but less frequent bursts in mutants compared to controls^7^. These findings in single neuron patch clamp experiments and those from network MEA recordings seem discordant.

To further probe this issue, we re-analyzed existing data from the same isogenic pairs in Mok et al. (2022)^7^. These data, starting at two weeks post-plating, used MEAs (Axion, 64-electrodes in 12-well plates) to record the network activity for 5 minutes every week for six weeks. Broadband time series were converted into digital events (spike rasters) representing the action potentials fired by neurons close to the recording electrodes. Here, we refer to the rate of action potentials across the different electrodes (i.e., channels) as network activity. As networks developed, the network activity transitioned through various stages. *MECP2* mutant and control networks developed spontaneous activity, first appearing as sparse firing two weeks post-plating (fig. S1a, b). By week 3, networks began synchronously bursting – forming network bursts. Network bursts first formed with a high degree of inter-burst spikes that ‘migrated’ into bursts over development (fig. S2). Strong synchronous bursting activity persisted until the end of the recording period.

In the example, the WIBR3 *MECP2* null networks at week four seem to develop longer bursts relative to the wildtype (WT). We zoomed into week six mutant and wildtype bursts and found that while the WT burst was composed of a single burst (fig. 2a), the mutant comprised a series of small, short bursts following an initial longer burst (fig. 2b).

**Figure 2.**
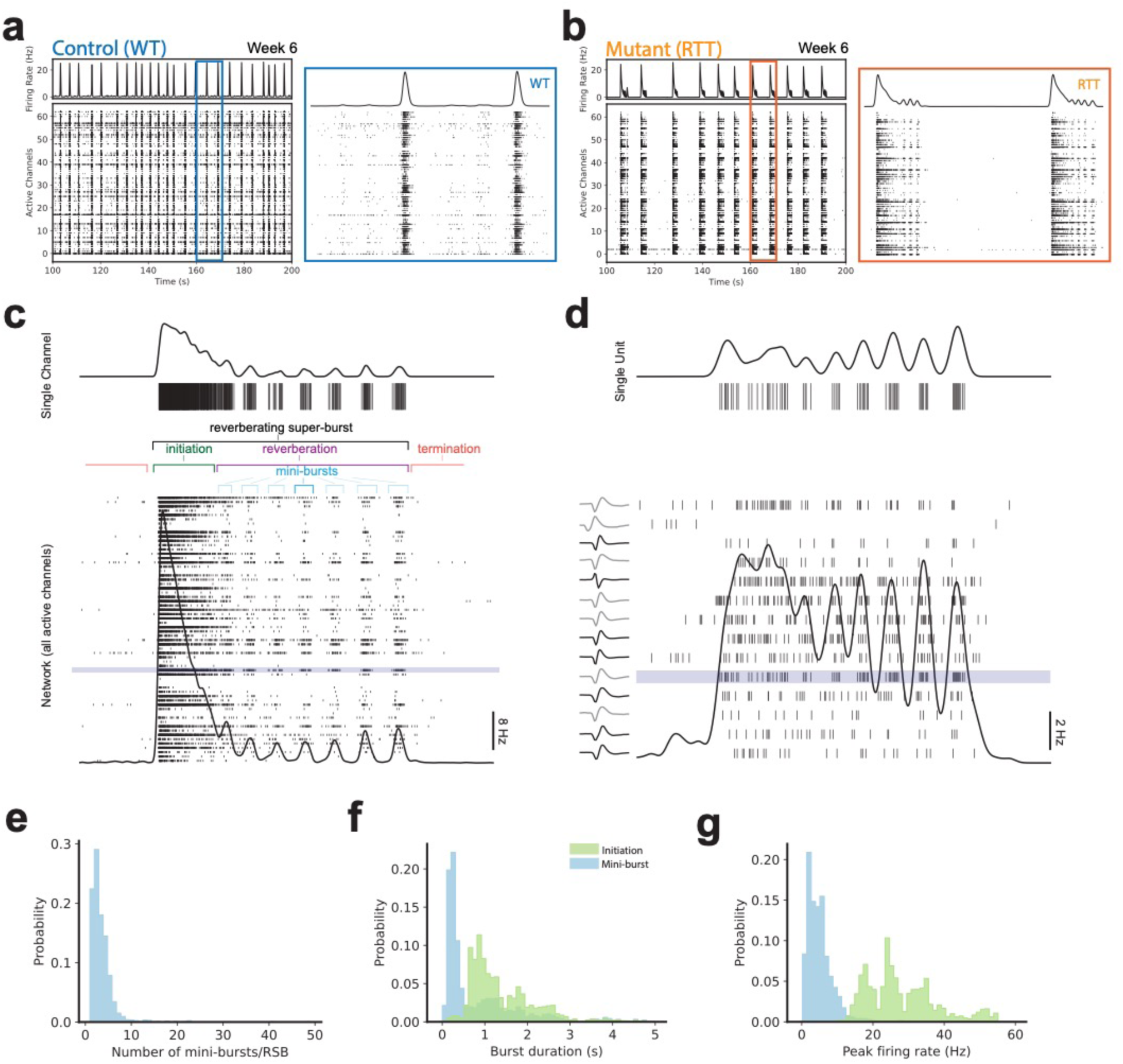
Reverberating super-bursts (RSBs) emerge in *MECP2* mutant networks and occur at a single unit, single channel, and network level. (a) Representative raster plot and associated spike density function of WIBR3 wildtype networks (6 weeks post-plating). (b) Representative raster plot and associated spike density function of WIBR3 null networks (6 weeks post-plating). (c) Bottom: Schematic of RSB, containing an initiation network burst followed by reverberations of multiple mini-bursts. Black trace represents spike density function. Top: Single channel activity highlighted from a selected channel (blue). (d) Bottom: Spike sorted activity of the same RSB seen in panel c). Top: Single unit activity highlighted from a selected unit (blue). Left: Extracellular action potential waveforms from each putative unit. (e) Distribution of the number of mini-bursts per RSB, (f) initiation burst (green) and mini-burst (blue) duration, and (g) initiation burst (green) and mini-burst (blue) peak burst firing rate.

### Reverberating super bursts (RSBs) emerge in MECP2 mutant neuronal networks

In a subset of wells of the WIBR3 null networks (fig. 2b), several network bursts emerged containing an initial single larger amplitude burst followed by several high-frequency lower amplitude bursts (fig. 2b, c). We refer to the former as the initiation network burst, and the succeeding lower amplitude bursts as mini-bursts (fig. 2c). The entire complex was termed Reverberating Super-Burst (RSB) due to a super burst structure^21^ that resembled acoustic reverberations (fig. 2b, c).

RSBs emerged at a network level, evident in the average instantaneous firing rate across all channels (fig. 2c *bottom*), but also appeared at a single channel (i.e., electrode) (fig. 2c *top*), and single unit (i.e., single neuron) level (fig. 2d). To demonstrate this, we built a spike density function (SDF) for each channel by convolving each spike with a bandwidth-optimized Gaussian kernel to approximate the instantaneous firing rate^30^. At the network level, the network averaged instantaneous firing rate demonstrated reverberating dynamics (fig. 2c). RSBs followed a stereotypical form (described in fig. 2c) with an initiation network burst with a high amplitude firing rate. Initiation network bursts had varying burst durations, usually below 500 milliseconds (fig. 2f), and a range of firing rates (fig. 2g). The succeeding mini-bursts varied in counts per RSBs (fig. 2e), duration (fig. 2f), and amplitude of firing rate (fig. 2g). RSBs ended with a period of quiescence that was longer than the interval between mini-bursts (fig. 2b, c). Thus, the initiation network burst was, in general, of longer duration (initiation burst median = 1.144s, mini-burst median = 0.359s) and larger amplitude (initiation burst median = 25.668Hz, mini-burst median = 3.891Hz) than the following mini-bursts. Interestingly, although mini-bursts were more frequent than initiation bursts, they show more narrow distributions of durations and firing rates, indicating that initiation bursts were more heterogeneous.

To investigate if RSBs were the product of multi-unit (i.e., multiple neurons) activity or single unit (i.e., single neuron) activity, we performed spike sorting using Plexon Offline Sorter in select reverberating wells (^31^, fig. 2d). In short, spike sorting attempts to decouple single-unit activity from multi-unit activity. After spike sorting, strict quality control for only putative single units resulted in most channels being discarded. Although most channels containing multi-unit activity could not be sorted and were therefore discarded, in the sorted channels, we show that RSBs persist at the single unit level (fig. 2d). However, RSB structure changed subtly, most notably in the amplitude of firing rate for the initiation network burst (fig. 2d). Thus, RSB is a phenomenon that also occurs in isolated single units as well as in multi-units. However, it also suggests complex ensemble dynamics underlying the origin of RSBs that cannot be extrapolated from single-unit recordings alone.

### Inconsistent handling and detection of RSBs

Network burst level features such as network burst frequency, and network burst duration are commonly reported as primary phenotyping metrics in many disease modelling studies^8^. In our *MECP2* mutant networks, RSBs exhibit various amplitude-frequency profiles (fig. S3). In order to examine how standard network burst detection algorithms handle reverberating activity, we re-analyzed the data using both fixed ISI (inter-spike-interval) and adaptive ISI network burst detection algorithms^32–34^. These approaches utilize a maximum ISI threshold that can distinguish spiking activity as occurring within versus outside of bursts – for example, ISIs smaller than the threshold would be classified as occurring within a burst, and vice versa^32,34^.

Fixed ISI network burst detection was highly susceptible to the experimenter’s choice of parameters compared to adaptive ISI network burst detection. Boundaries around bursts (i.e., the start and end of bursts) were sensitive to parameter choice and burst detection method (fig. S4). Differences in burst boundaries produced significant differences in burst features such as network burst frequency (One-way ANOVA, *F*(2,41) = 23.034, *p* < 0.001, fig. 3a top) and network burst duration (One-way ANOVA, *F*(2,17.048) = 12.640, *p* < 0.001, fig. 3a bottom). Interestingly, post hoc comparison for network burst frequency revealed that adaptive ISI burst detection was not significantly different compared to 100ms ISI fixed-based burst detection (*p_bonf_* = 0.585), whereas 15ms ISI fixed-based burst detection was significantly different from both adaptive and 100ms ISI fixed-based methods (*p_bonf_* < 0.001, *p_bonf_* < 0.001, respectively). On the other hand, post hoc comparison for network burst duration revealed our choice of adaptive-based burst detection was significantly different compared to 100ms ISI fixed-based burst detection (*p_bonf_* = 0.024) but not 15ms ISI fixed-based burst detection (*p_bonf_* = 0.079). Again, 15ms ISI fixed-based burst detection significantly differed from the 100ms ISI fixed-based burst detection (*p_bonf_* < 0.001). Based on this, the adaptive ISI burst detection method determines burst boundaries variably either around the entire RSBs or the individual mini-bursts, which would therefore impact the reliability of downstream bursting metrics. We also conducted a similar Welch power spectral density (PSD) estimate seen in Mok et al. (2022)^7^. This approach enables the characterization of the frequency components contained in the network’s spike trains. PSD detected the burst frequency in both reverberating and non-reverberating networks (fig. 3b *red dot*); however, could not estimate the faster reverberation frequency of mini-bursts (fig. 3b *bottom red arrow*).

**Figure 3.**
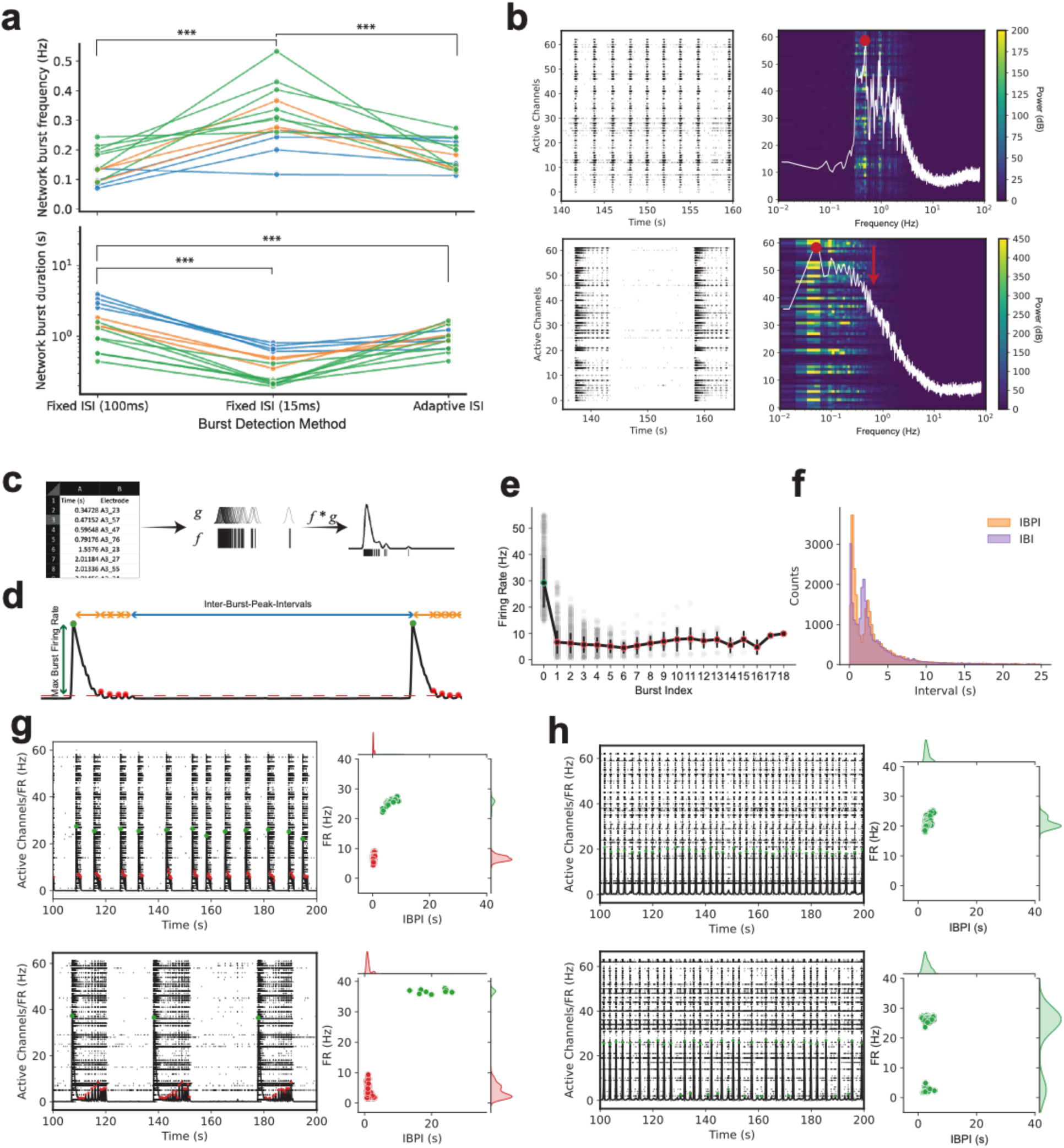
Inconsistent handling by standard burst detection algorithms and two-feature detection of reverberating networks. (a) Comparison of network burst frequency (top), and network burst duration (bottom) by different ISI-based burst detection algorithms. Colours indicate replicates from the same plate. (b) Burst detection using power spectral density (PSD) estimates (previously used in Mok et al., 2022). Non-reverberating networks (top) present with a sharp peak at the network burst frequency (red dot). Reverberating networks (bottom) present with a broad peak at network burst frequency (red dot) but no peak at expected mini-burst frequency (red arrow). White traces represent the average of the PSD for the network. (c) Calculation of spike density function (SDF) by convolution of spike time with optimized Gaussian kernel. SDF represents an estimate of the instantaneous firing rate of the network. (d) Schematic of burst peak features. Bursts were identified by finding local peaks that exceed a minimum burst firing rate threshold (height) and change in firing rate (prominence). Dots represent detected burst peaks. (e) Burst peak firing rate (amplitude) relative to burst position (index) within an RSB. A burst position of 0 represents the initiation burst. Burst positions greater than 0 represent the mini-bursts. Initiation burst has a larger amplitude than mini-bursts. Burst amplitude can be used to label bursts as either initiation bursts or mini-bursts. (f) Distribution of inter-burst-peak-interval (IBPI) approximates the distribution of inter-burst-interval (IBI). IBI is left-shifted relative to IBPI. (g) Examples of clustering results for reverberating networks. (h) Examples of clustering results for non-reverberating networks. Error bars (in f) represent standard deviation. Red dots (in g-h) represent labelled mini-bursts, whereas green dots represent initiation network bursts. Statistical significance was evaluated using One-Way ANOVA. \**p* < 0.05, \*\**p* < 0.01, \*\*\**p* < 0.001.

The consistent and accurate detection of network bursts by automated detection algorithms commonly used in high-throughput analyses is essential for phenotypes to be reproducible. The aforementioned burst detection methods capture and describe the same phenomena in different, incomplete ways. We found that the experimenter’s choice of burst detection algorithm and parameters significantly impacted where network burst boundaries were drawn within RSBs, which would therefore impact the accuracy of network phenotypes in subsequent downstream analyses.

To develop an algorithm that reliably identifies RSBs, we first operationally defined an RSB. An RSB is an activity pattern that contains various repetitive low-amplitude and high-frequency mini-bursts nested within a larger network event. Specifically, a large amplitude network burst must precede the smaller amplitude mini-bursts (fig. 3d, e). The inter-burst-intervals (IBIs) within the larger network event (typically of the mini-bursts) must be shorter than the IBIs between subsequent large network events, here the RSBs (fig. 3d, f).

Using the network-level SDF, we identified local activity peaks that met a criterion (see Materials and Methods, fig. 3d). Local activity peaks mapped onto network bursts with a sufficient firing rate (fig. 3d, e). Inter-burst-peak-interval (IBPI), the time between subsequent local activity peaks, approximated IBI. IBPI was used in reverberating network detection because it offered an adequate approximation of IBI that was unbiased by the calculation of the burst’s boundaries (fig. 3f). This approach is useful in the detection of RSBs because burst boundary detection is non-trivial when the network burst firing rate does not reach an arbitrarily low threshold before increasing again – the case in some RSBs (fig. 3d). The amplitude of the peak represented the maximum firing rate of each network burst (fig. 3d). Amplitude was used to label bursts as either initiation network bursts or mini-bursts (fig. 3d, e). IBPI was used to determine when bursts were part of or outside of RSBs (see R_max_ calculation under *Reverberating Super Burst Construction Loop* in Materials and Methods).

Reverberating networks were identified by plotting the amplitude of the detected bursts against their frequency (IBPI). Reverberating networks show two clear clusters (fig. 3g). The first cluster was found to populate the high firing rate and long IBPI dimensions. This represented the large amplitude initiation network burst and the corresponding long interval to the mini-burst of the previous RSB (fig. 3g *green dots in the scatter plot*). The second cluster populated to low firing rate mini-bursts and low-to-medium IBPI to the previous mini-burst or to the large initiation burst within a RSB (fig. 3g *red dots in the scatter plot*).

In non-reverberating networks, our analyses often produced a single continuous cluster (fig. 3h *green dots in the top scatter plot*). The single cluster was tightly constrained to a narrow range, representing a highly periodic network with regular IBPIs and a similar maximum firing rate within each network burst. Occasionally, in some networks, where the amplitude of the maximum firing rate for each network burst was highly variable, two clusters would appear but overlap strongly along the IPBI dimension (fig. 3h *green dots in the bottom scatter plot*). Since the frequency of mini-bursts is essential to reverberations we only considered reverberating networks when clusters could be distinguished along the IPBI dimension or along both dimensions (fig. 3g). The latter was an appropriate method since we rarely found clusters that segregated along the IBPI dimension but not along the firing rate dimension.

This approach allowed us to categorize networks as reverberating or non-reverberating. Furthermore, labels and parameters were extracted to identify the types of bursts (e.g., initiation network burst vs. mini-burst) and the threshold to adaptively define boundaries around RSBs for subsequent feature calculation (see R_max_ calculation under *Reverberating Super Burst Construction Loop* in Materials and Methods and fig. S5 for algorithm block diagram).

### MECP2 null networks reverberate more than isogenic control networks

We categorized network activity into three groups based on the proportion of RSBs relative to the total number of detected bursts in the three isogenic *MECP2* mutant and control pairs. These activity groups are: 1) Non-reverberating networks (containing less than 20% RSBs), 2) Mixed-reverberating networks (containing between 20%-70% RSBs), and 3) Reverberating networks (containing more than 70% RSBs). Comparisons of the distributions of reverberating networks between the isogenic pairs show more RSBs in WIBR3 and PGPC14 *MECP2* null networks at different development stages (fig. 4a). RSBs were detected to equivalent levels in the CLT pairs. We introduced the Network Reverberation Index (NRI) metric to describe this variable distribution (see NRI Calculation in Materials and Methods). A positive NRI means that the *MECP2* mutant network contains more RSBs than isogenic controls. A negative NRI means the opposite and a zero NRI means they contain the same number of RSBs.

**Figure 4.**
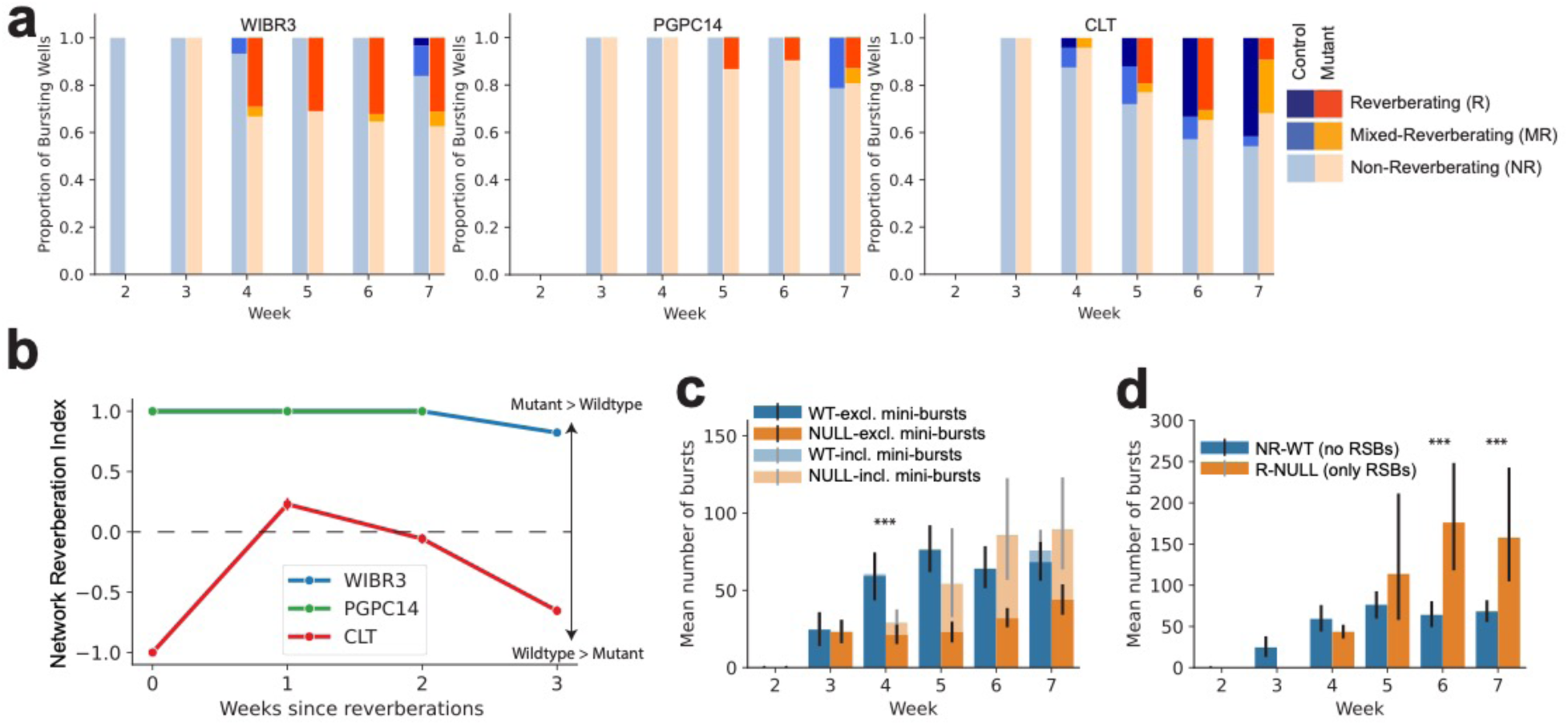
*MECP2* null networks reverberate more than isogenic controls and exhibit greater bursting activity, indicating hyper-activity. (a) Proportion of reverberating, mixed-reverberation, and non-reverberation wells relative to the total number of bursting wells across development for each individual. (b) Network Reverberation Index, representing the plate-matched difference in proportion of reverberating networks for each individual. (c) Mean number of network bursts per recording when mini-bursts are included (incl.) and excluded (excl.) in all WIBR3 control (WT) and null (NULL) networks. (d) Mean number of network bursts per recording for non-reverberating control (NR-WT) and reverberating null (R-NULL) networks. All bar plots use mean +/− 95% confidence intervals. Statistical significance was evaluated using Kruskal-Wallis test and Two-Way ANOVA. \**p* < 0.05, \*\**p* < 0.01, \*\*\**p* < 0.001.

We show that WIBR3 null networks consistently show higher NRI values relative to isogenic controls networks (fig. 4a *left,* b) during the middle (weeks 4 and 5) to late (week 6 and 7) stages of development (Kruskal-Wallis, *H*(1) = 16.653, *p* < 0.001, fig. 4a left). For PGPC14 null networks, the NRI values also showed a higher proportion of reverberating networks compared to their controls (fig. 4b) but these were not significantly different from isogenic pairs (Kruskal-Wallis, *H*(1) = 0.419, *p* = 0.517) due to the low numbers of reverberating networks (fig. 4a middle). The NRI values close to zero or less for the CLT mutant networks showed no significant differences in the proportion of RSBs relative to isogenic controls (Kruskal-Wallis, *H*(1) = 2.344, *p* = 0.126, fig. 4a *right,* b). This pattern seems consistent with the milder atypical Rett Syndrome phenotype reported in this patient by Mok et al. (2022)^7^.

Importantly, when considering mini-bursts contributing to the total number of network bursts WIBR3 null networks show increased network bursts relative to controls (log-transformed Two-way ANOVA, *F*(3,167) = 3.888, *p* = 0.010, fig. 4d). Post hoc pairwise comparison between WIBR3 wildtype controls and null groups at specific developmental time points revealed the number of network bursts was significantly increased relative to isogenic controls during week 6 (*p_bonf_* < 0.001) and 7 (*p_bonf_* = 0.018) (fig. 4d).

Thus, our results portray a different picture than in the previous study^7^. *MECP2* null neurons assemble into networks that produce a larger proportion of RSBs than isogenic controls. The increase in RSB causes an increase in the total number of network bursts and therefore network burst frequency, a trend that becomes more obvious during weeks 6 and 7, which could be considered the latter developmental stages of these networks (fig. 4d). This result indicates *MECP2* null mutants networks are more excitable than isogenic controls, which corresponds to the more excitable single neurons phenotypes reported in Mok et al. (2022)^7^.

### RSBs bias the calculation of burst metrics

To further investigate RSBs in *MECP2* mutants and controls, we focused on the WIBR3 networks where the RSB phenotype was most pronounced. We first identified WIBR3 networks that exhibited RSBs and stratified the replicates into four groups: non-reverberating control (NR-WT) networks, non-reverberating *MECP2* null (NR-NULL) networks, reverberating control (R-WT) networks, and reverberating null (R-NULL) networks (fig. 5a). These categories were based on the previously stated proportion (i.e., reverberating networks containing greater than 70% RSBs, non-reverberating networks containing less than 20% RSBs, and excluding mixed-reverberating networks which contain between 20%-70% RSBs).

**Figure 5.**
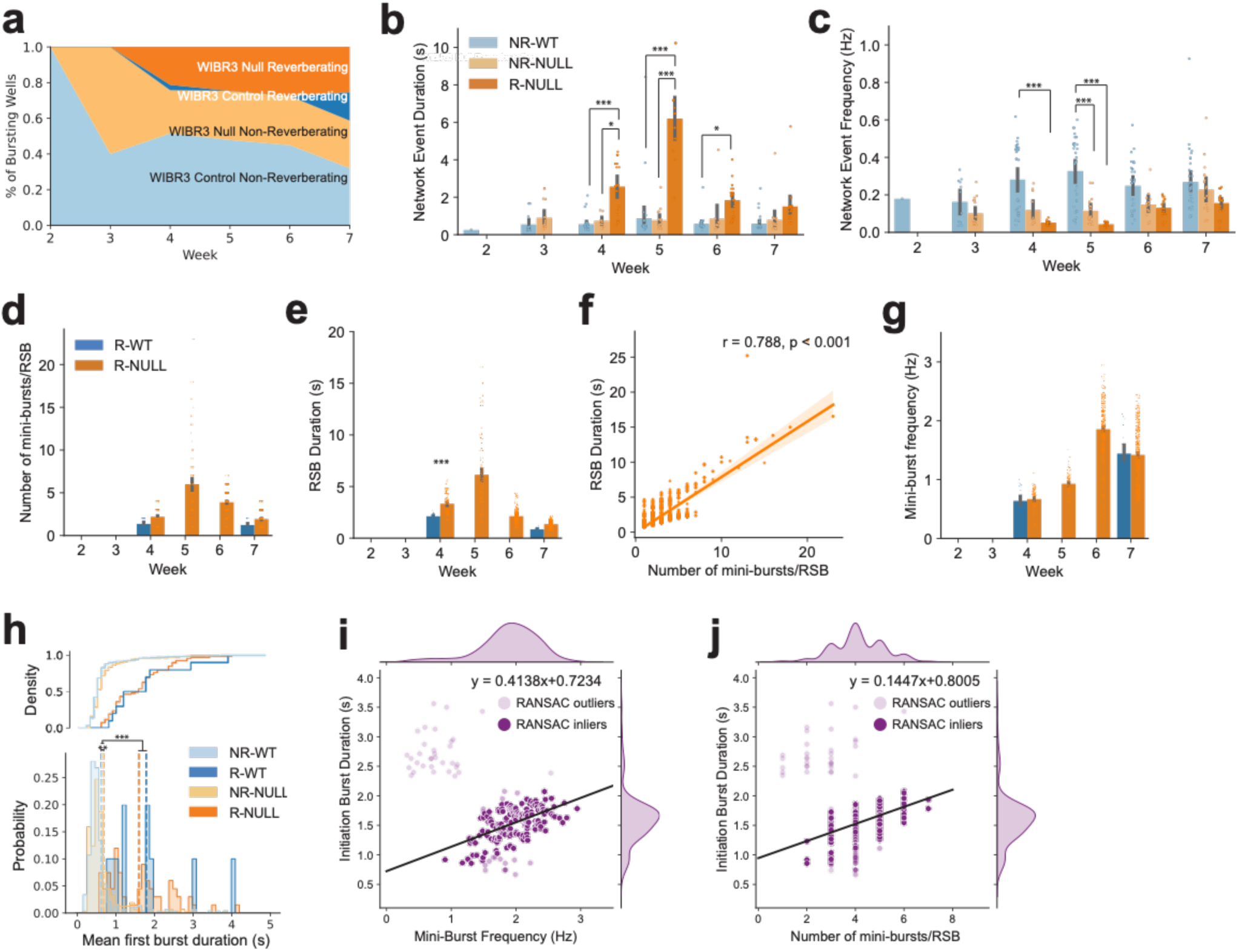
RSBs alter network bursting metrics. (a) Wildtype and *MECP2* null WIBR3 networks were stratified as either non-reverberating or reverberating. (b and c) Quantification of (b) network event duration and (c) network event frequency between non-reverberating control networks (NR-WT), non-reverberating mutant networks (NR-NULL), and reverberating mutant networks (R-NULL). Network event refers to the entire RSBs (initiation network burst and mini-bursts) in reverberating networks or discrete single network bursts in non-reverberating networks. (d) Quantification of the number of mini-bursts per RSB of reverberating WIBR3 WT (R-WT) and WIBR3 null (R-NULL) networks. (e) Quantification of RSB duration of reverberating R-WT and R-NULL networks. (f) Linear regression of number of mini-bursts per RSB and RSB duration in R-NULL networks. Suggests the long network burst duration is associated with an increased number of mini-bursts that were unaccounted for. (g) Quantification of reverberating mini-burst frequency of R-WT and R-NULL networks. (h) Distribution of initiation burst duration in reverberating (R-WT and R-NULL) networks and discrete single network burst duration of non-reverberating (NR-WT and NR-NULL) networks. (i) RANSAC regression of mini-burst frequency and initiation burst duration in week 6 R-NULL networks. Suggests longer initiation burst duration is associated with increased reverberating mini-burst frequency. Lighter colored points indicate RANSAC outliers. Darker colored points indicate RANSAC inliers. Linear fit performed on RANSAC inliers. See RANSAC (RANdom Sample Conensus) Regression under Materials and Methods. (j) RANSAC regression of number of mini-bursts per RSB and initiation burst duration in week 6 R-NULL networks. Suggests longer initiation burst duration is associated with increased number of mini-bursts that follow. Lighter colored points indicate RANSAC outliers. Darker colored points indicate RANSAC inliers. Linear fit performed on RANSAC inliers. See RANSAC (RANdom Sample Conensus) Regression under Materials and Methods. NR: Non-reverberating. R: Reverberating. All bar plots use mean +/− 95% confidence intervals. Statistical significance was evaluated using Two-way ANOVA, Kruskal-Wallis Test, Linear regression, and RANSAC regression. \**p* < 0.05, \*\**p* < 0.01, \*\*\**p* < 0.001.

In the following comparisons, a network event refers to the entire RSBs (initiation network burst and mini-bursts) in reverberating networks or discrete single network bursts in non-reverberating networks. This was done to establish a link to the method used in Mok et al. (2022) that documented a decrease in network bursts in WIBR3 null mutants relative to controls (fig. S6)^7^. Network event duration was significantly different between WIBR3 NR-WT, NR-NULL, and R-NULL groups (Two-way ANOVA, *F*(2,250) = 72.051, *p* < 0.001, fig. 5b) and between different developmental time points (Two-way ANOVA, *F*(4,250) = 22.577, *p* < 0.001, fig. 5b), with a significant interaction (Two-way ANOVA, *F*(8,250) = 13.983, *p* < 0.001). Post-hoc comparison revealed burst duration in NR-WT and NR-NULL were not significantly different (*p_bonf_* = 0.788), whereas R-NULL was significantly different from NR-NULL (*p_bonf_* < 0.001) and NR-WT (*p_bonf_* < 0.001). Thus, network event duration was longer in R-NULL, which would be anticipated if such events are RSBs.

Network event frequency was significantly different between NR-WT, NR-NULL, and R-NULL groups (Two-way ANOVA, *F*(2,252) = 34.095, *p* < 0.001, fig. 5c) and between different developmental time points (Two-way ANOVA, *F*(4,252) = 4.464, *p* < 0.01, fig. 5c), with a non-significant interaction (Two-way ANOVA, *F*(8,252) = 1.674, *p* = 0.105, fig. 5c). Post-hoc comparison revealed all groups were statistically different from each other (NR-NULL*R-NULL: *p_bonf_* < 0.05; NR-NULL*NR-WT: *p_bonf_* < 0.001; R-NULL*NR-WT: *p_bonf_* < 0.001). The NR-NULL networks had significantly lower network event frequency compared to the NR-WT networks, unlike network event duration which did not show a significant difference (fig. 5b-c). However, the lowest network event frequency was in the R-NULL networks (fig. 5c). The latter would be also anticipated in networks that show an increase in RSBs.

To quantitatively explore the factors that influenced the changes in RSB duration and RSB frequency in reverberating networks, we examined the reverberating mini-burst activity in both reverberating WIBR3 wildtype (R-WT) and null (R-NULL) networks that only had RSB network events. We found that as the R-NULL networks developed, they consistently had elevated numbers of mini-bursts per RSBs compared to the R-WT networks (Kruskal-Wallis, *H*(1) = 35.126, *p* < 0.001, fig. 5d). The peak of reverberating activity occurred during week 5. Similarly, R-NULL networks have significantly longer RSB duration relative to controls (Kruskal-Wallis, *H*(1) = 451.899, *p* < 0.001, fig. 5e). Through linear regression, we found that the longer RSBs duration in all R-NULL networks was positively correlated with an increased number of mini-bursts per RSBs (Pearson correlation coefficient r = 0.788, p < 0.001, fig. 5f).

Furthermore, we calculated the mini-burst frequency within each RSB. We found that mini-burst frequency was significantly elevated in R-NULL compared to isogenic R-WT networks (Kruskal-Wallis, *H*(1) = 10.204, *p* = 0.001). The mini-burst frequency peaked during week 6 and subsequently decreased during week 7 (fig. 5g). Taken together, that change in RSB duration (fig. 5e), which decreased by 2-fold in R-NULL networks after week 5, was likely due to decreased number of mini-bursts per RSB (fig. 5d) and increased mini-burst frequency (fig. 5g). Importantly, these results show that even if RSB were not exclusively found in R-NULL networks, their features were different from those found in R-WT networks suggesting that MECP2 deficiency exacerbates the RSB phenotype.

Most RSBs initiate with a longer-than-normal network burst. It is possible that the length of the initiation burst determines the probability that mini-bursts will follow. We compared the first initiation burst duration in reverberating (R) networks to non-reverberating (NR) discrete single network burst duration. Duration of the initiation burst was significantly different from burst duration between reverberating and non-reverberating networks (Kruskal-Wallis, *H*(3) = 4393.402, *p* < 0.001, fig. 5h). Post hoc pairwise comparisons showed that reverberating control (R-WT) and reverberating null (R-NULL) networks had significantly increased first burst duration compared to burst duration of non-reverberating control (R-WT) and non-reverberating null (R-NULL) networks (fig. 5h). Finally, we found that longer initiation burst duration is associated with a faster mini-burst frequency (RANSAC linear fit of week 6 inliers, m = 0.4138, fig. 5i) and greater number of mini-bursts per RSB (RANSAC linear fit of week 6 inliers, m = 0.1447, fig. 5j). These findings reflect the idiosyncratic developmental trajectory of *MECP2* null compared to isogenic controls.

### GABA antagonism does not affect RSBs

A previous study reported that reverberations in stem cell-derived organoids were sensitive to the application of Bicuculline, a GABA antagonist^26^. This seems at odds with our results since our monolayer networks were composed of excitatory neurons^7^. To investigate this issue, we performed new MEA studies on the WIBR3 excitatory networks, since these were the ones showing the stronger RSB phenotype. Notably, in this separate experiment, we validated the previously discussed skew of RSBs towards the null networks, with very few control networks exhibiting RSBs.

Application of bicuculline in non-reverberating control (NR-WT) and reverberating null (R-NULL) excitatory networks had no effect on the presence of RSBs (NR-WT: Wilcoxon Signed-Rank, *Z* = –0.365, *p* = 0.855; R-NULL: Paired Student T, t(15) = 1.463, *p* = 0.164, fig. 6a-b). Interestingly, in both bicuculline-treated networks, there was a subtle significant increase in network burst frequency (NR-WT: Paired Student T, t(17) = –3.700, *p* = 0.002; R-NULL: Wilcoxon signed-rank, *Z* = –3.103, *p* = 0.001, fig. 6c) but not network burst duration (NR-WT: Wilcoxon signed-rank, *Z* = 0.849, *p* = 0.417; R-NULL: Paired Student T, t(15) = 1.604, *p* = 0.130, fig. 6d). The former may be due to off-target effects of bicuculline on membrane channels such as small conductance calcium-activated potassium channels which is responsible for the slow afterhyperpolarization of neurons after action potential firing ^35,36^. Importantly, no significant differences were found between pre– and post-bicuculline-treated R-NULL networks in mini-burst frequency (Paired Student T, t(15) = –0.639, *p* = 0.532, fig. 6e). These results corroborate our networks were predominantly excitatory, and bicuculline had no effect on the RSB phenotype.

**Figure 6.**
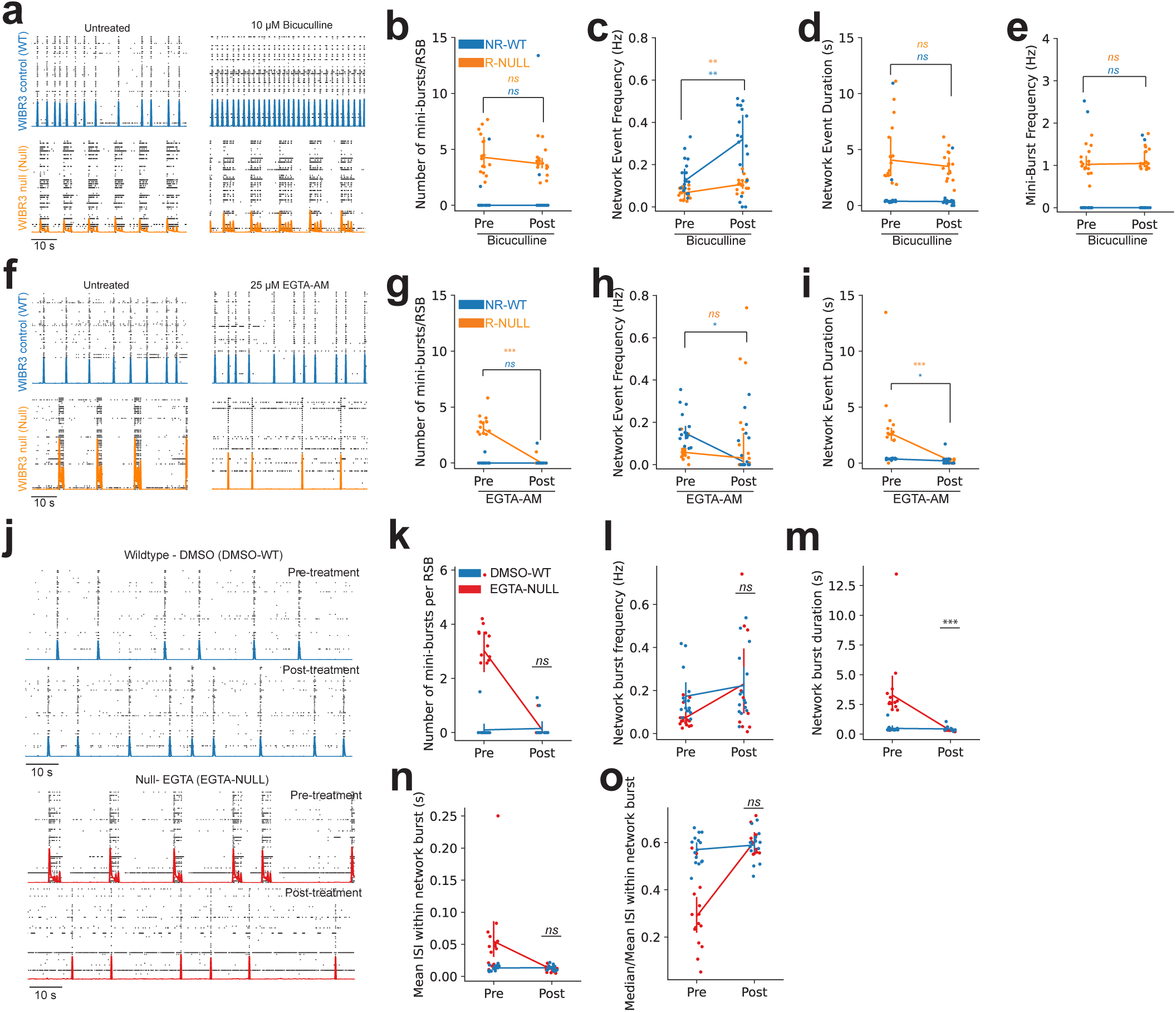
Pharmacological treatment of EGTA-AM and not bicuculline eliminates RSBs and rescues network bursting phenotype. (a) Representative raster plot of WIBR3 wildtype (WT) and null (NULL) networks before and after bicuculline treatment. WIBR3 WT networks did not reverberate (NR-WT) whereas WIBR3 null networks did (R-NULL). (b-e) Quantification of (b) Number of mini-bursts per RSB, (c) Network event duration, (d) Network event frequency), and (e) Mini-burst frequency in pre– and post-bicuculline treated WIBR3 networks. (f) Representative raster plot of WIBR3 NR-WT and R-NULL networks before and after EGTA-AM treatment. (g-i) Quantification of (g) Number of mini-bursts per RSB, (h) Network event duration, and (i) Network event frequency) in pre– and post-EGTA-AM treated WIBR3 networks. (j) Top: Representative raster plot of WIBR3 NR-WT networks before and after DMSO treatment. Bottom: Representative raster plot of WIBR3 R-NULL networks before and after EGTA treatment. (k-o) Quantification of (k) Number of mini-bursts per RSB, (l) Network burst frequency, (m) Network burst duration, (n) Mean ISI within network bursts, and (o) Median/Mean ISI within network bursts in pre– and post-DMSO treated WT networks (DMSO-WT) and pre– and post-EGTA-AM treated NULL (EGTA-NULL) networks. Point plots use mean (b-e and g-i) and median (k-o) +/− 95% confidence intervals. Statistical significance was evaluated using Mann-Whitney test, Wilcoxon Matched Pairs Signed Rank test, Independent Samples Student T-test, and Paired Student T-test. \**p* < 0.05, \*\**p* < 0.01, \*\*\**p* < 0.001.

### RSBs are dependent on asynchronous Ca^2+^ release

We hypothesized that the mechanisms responsible for RSBs may operate at the level of the excitatory synapse. So, it is possible that by down-regulating the amount of neurotransmitter release by partially blocking Ca^2+^ availability in the presynaptic terminal, the RSB phenotype could be rescued. We applied ethylene glycol tetraacetic acid acetoxymethyl ester (EGTA-AM), which decreases Ca^2+^ and vesicular neurotransmitter release in the microdomain of single channels when they are not saturated^37^. We reasoned that during low-amplitude mini-bursts, channels might not be saturated, and EGTA-AM may bind to Ca^2+^ and decrease its availability at the presynaptic vesicular sensor. The latter would decrease the frequency of mini-bursts. On the other hand, during typical high-amplitude bursts, Ca^2+^ would be more abundant, and channels may be saturated. In the latter scenario, EGTA-AM would have none or a small effect on network bursts.

EGTA-AM was applied to both non-reverberating control (NR-WT) and reverberating null (R-NULL) networks (fig. 6f). EGTA-AM had a significant effect on the presence of RSBs (Wilcoxon signed-rank, *Z* = 3.237, *p* = 0.001, fig. 6g). EGTA-AM robustly eliminated RSBs in R-NULL networks (Paired Student T, t(15) = 6.722, *p* < 0.001, fig. 6f, g) and had no effect on RSBs in NR-WT networks (Wilcoxon signed-rank, *Z* = –0.447, *p* = 1.000, fig. 6f, g). Furthermore, EGTA-AM had no significant effect on network event frequency in R-NULL networks (Wilcoxon signed-rank, *Z =* –0.362, *p* = 0.737, fig. 6h). However, as RSBs and therefore the mini-bursts were eliminated, EGTA-AM significantly decreased network event duration (Paired Student T, t(15) = 5.566, *p* < 0.001, fig. 6i) in previously R-NULL networks but also had a significant effect in NR-WT networks (Wilcoxon signed-rank, *Z* = 2.499, *p* = 0.010, fig. 6i).

We further compared EGTA-AM treated null network (EGTA-NULL, fig. 6j *bottom*) features to dimethyl sulfoxide (DMSO) vehicle-treated control network (DMSO-WT, fig. 6j *top*) features. This provided an additional control for the possible effects of experimental manipulations on the network activity. EGTA-AM treatment eliminated RSB in mutant networks. Remarkably, we found no significant differences in the presence of RSBs between post-treatment EGTA-NULL and DMSO-WT groups (Mann-Whitney, *p* = 0.635, fig. 6k). Network burst frequency was not significantly different between post-treatment EGTA-NULL and DMSO-WT (Mann-Whitney, *p* = 0.756, fig. 6l). However, burst duration was significantly different between post-treatment EGTA-NULL and DMSO-WT (Mann-Whitney, *p* < 0.001, fig. 6m) with DMSO-WT having longer average burst duration (Mean burst duration, DMSO-WT = 0.436s, EGTA-NULL = 0.292s).

Both mean ISI within network bursts (Paired Student T, t(23) = –1.522, *p* = 0.142, fig. 6n) and median/mean ISI within network bursts (Paired Student T, t(23) = 0.598, *p* = 0.555, fig. 6o) were not significantly different between post-treatment groups (EGTA-NULL and DMSO-WT), indicating both networks had similar firing rate distributions (as ISI is the inverse of firing rate). In summary, EGTA-AM treatment of reverberating null networks reduces RSBs while preserving network bursts with similar intra– and inter-bursting dynamics to non-reverberating WT networks.

### EGTA-AM decreases the duration of the initiation network burst

As RSBs initiate with a longer-than-normal network burst, one may ask whether EGTA-AM also reduces the duration of the first network burst. It is possible that the rescue by EGTA-AM was at the expense of a reduction in the duration of the initial network burst. To test this hypothesis, we compared the burst duration of EGTA-treated null (EGTA-NULL) networks to DMSO vehicle-treated isogenic wildtype control (DMSO-WT) networks.

DMSO vehicle treatment had no significant effect on the duration of discrete network bursts in non-reverberating control (DMSO-WT) networks or the initiation burst duration in reverberating null (DMSO-NULL) networks (Kruskal-Wallis, Dunn’s post hoc, *p_bonf_* = 1.000 and *p_bonf_* = 1.000, respectively, fig. 7a). On the other hand, EGTA-AM reduced the duration of the initiation burst in previously reverberating null (EGTA-NULL) networks (Kruskal-Wallis, Dunn’s post hoc, *p_bonf_* < 0.001, fig. 7b-d). Network burst duration in EGTA treated null (EGTA-NULL) networks was significantly shorter than control networks without EGTA-AM (preEGTA-WT) treatment (Kruskal-Wallis, Dunn’s post hoc, *p_bonf_* = 0.026) but not significantly different from WT with EGTA (EGTA-WT) networks (Kruskal-Wallis, Dunn’s post hoc, *p_bonf_* = 0.663, fig. 7b). Furthermore, EGTA-WT and preEGTA-WT were not significantly different from each other (Kruskal-Wallis, Dunn’s post hoc, *p_bonf_* = 0.456, fig. 7b). These results are more easily seen in Figure 7c, where the SDFs for the burst of pre– and post-EGTA treatment in control and nulls are overlaid on top of each other. A stark difference can be seen when comparing EGTA-NULL and preEGTA-NULL. Along the same lines, we found that the initiation burst duration was significantly reduced in EGTA-NULL compared to preDMSO-WT (Kruskal-Wallis, Dunn’s post hoc, *p_bonf_* = 0.011, fig. 7d) and DMSO-WT (Kruskal-Wallis, Dunn’s post hoc, *p_bonf_* = 0.031, fig. 7d).

**Figure 7:**
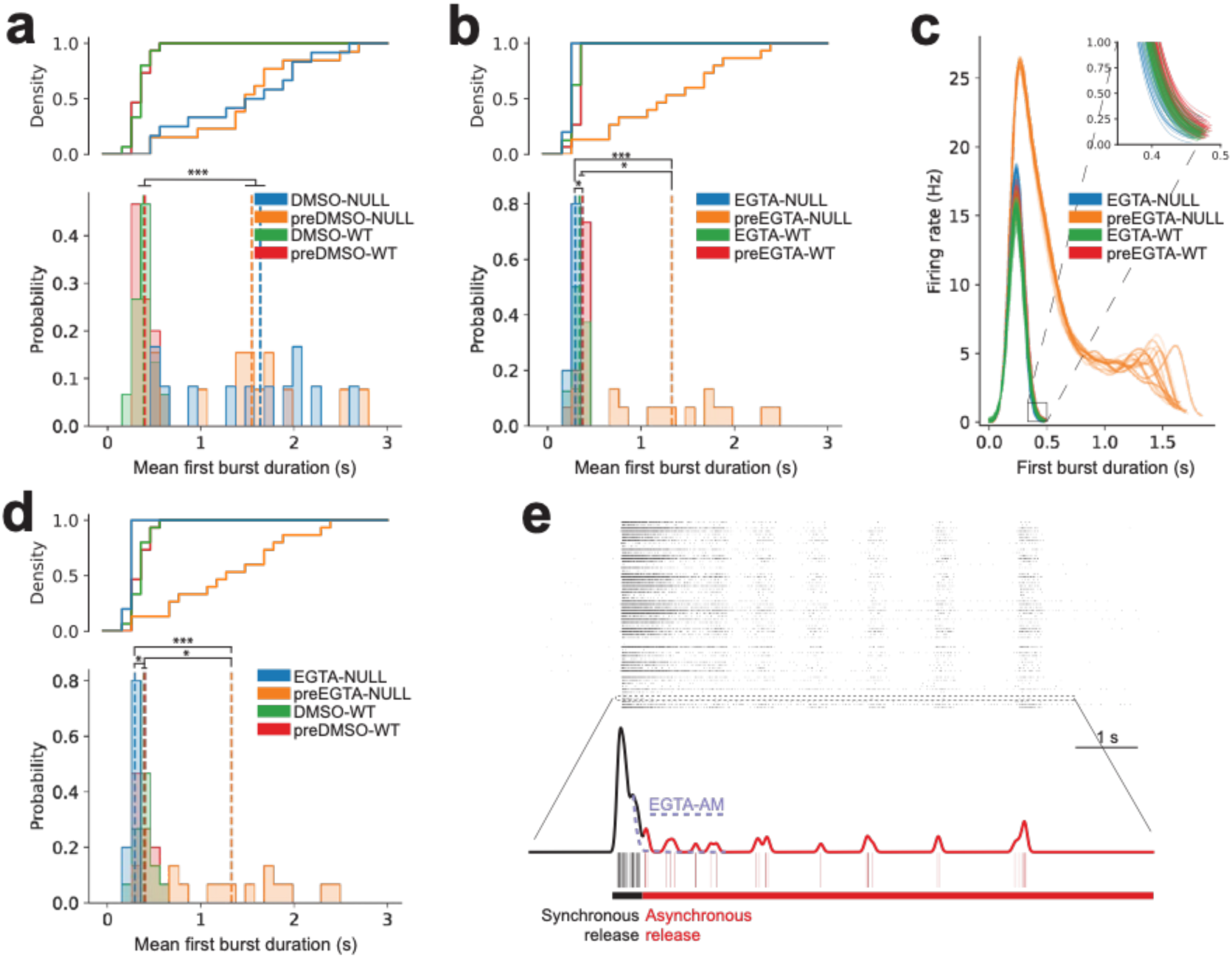
EGTA-AM reduces the duration of the initiation network burst in RSBs. (a, b, d) Cumulative distribution function (CDF) and associated histogram of mean burst duration of non-reverberating wildtype (WT) networks compared to the duration of the initiation network burst of reverberating *MECP2* null networks (NULL). Dashed lines represent the mean burst duration per group. (a) Distribution of burst duration in response to DMSO treatment to both NULL (DMSO-NULL) and WT (DMSO-WT). (b) Distribution of burst duration in response to EGTA treatment to both NULL (EGTA-NULL) and WT (EGTA-WT). (c) Overlaid truncated SDF of each the first burst duration for preEGTA-WT, preEGTA-NULL, EGTA-WT, and EGTA-NULL. Inset shows subtle differences between first burst duration in reverberating group (preEGTA-NULL) and non-reverberating groups (preDMSO-WT, DMSO-WT, EGTA-NULL). Representative networks were used. (d) Distribution of burst duration in response to DMSO treatment in WT (DMSO-WT) networks and EGTA treatment in NULL (EGTA-NULL) networks. (e) Hypothesized mechanism for elongated initiation burst in RSBs. The dashed line represents the effect of EGTA-AM treatment on initiation burst duration. Statistical significance was evaluated using One-Way ANOVA, Kruskal-Wallis Test, and Dunn’s post hoc comparisons. \**p* < 0.05, \*\**p* < 0.01, \*\*\**p* < 0.001.

These results suggest that the long duration of the first network burst may be triggering the mini-bursts and that by reducing the availability of Ca^2+^, and therefore asynchronous neurotransmitter release the hyperexcitable reverberating phenotype can be rescued (fig. 7e). In the discussion, we will speculate on possible mechanisms for this effect.

## DISCUSSION

In the current study, we investigated the development of *MECP2* mutant *Ngn2* human PSC derived excitatory networks using electrophysiological methods and compared them to isogenic controls. We report the following findings: (1) *MECP2* null neurons show an increase in excitability and spike frequency adaptation in intracellular recordings relative to isogenic controls, (2) *MECP2* null networks explored using MEA recordings show an increase in the proportion of RSB consisting of a large initial network burst followed by several smaller amplitude high-frequency mini-bursts that overall produce greater network burst frequencies relative to isogenic controls, and (3) administration of EGTA-AM (a slow-kinetic membrane permeable Ca^2+^ chelator) decreases RSBs rescuing the bursting network phenotype relative to isogenic controls.

### Single neuron and network excitability in MECP2 null neurons

Previous studies have shown that single neurons with *MECP2* null mutations show smaller cell bodies with decreased dendritic branching relative to isogenic controls as well as increased input resistance, decreased capacitance, and reduced Na^+^ and K^+^ currents^7^. Here, we corroborate these reports and showed that *MECP2* null neurons have indeed reduced rheobase – the amount of current necessary to evoke action potentials using current-clamp recordings^7^. Moreover, we found that spike frequency adaptation of spike trains evoked by a 1-second-square current pulse was increased in *MECP2* null relative to isogenic controls neurons. A decrease in action potential amplitude between subsequent spikes also appeared, further substantiating the decreased Na^+^ currents reported in Mok et al. (2022). These findings can be interpreted as *MECP2* null neurons showing an increased excitability accounting for the decrease in rheobase to elicit action potentials as a function of stimulation intensity and also increased adaptation which accounts for the fast decay in firing rate during sustained stimulation.

A somewhat contradictory finding was that despite this hyperexcitability in single neurons, network burst frequency in *MECP2* null networks was originally determined by standard burst detection algorithms to be decreased^7^. Specifically, the *MECP2* null networks showed a decreased number of network bursts, increased network burst duration, and decreased network burst frequency^7^. These findings were interpreted as a decrease in network excitability. But how can hyperexcitable *MECP2* null neurons produce a hypoexcitable phenotype? Here, we zoomed into the long bursts of *MECP2* null neurons and found a phenomenon we termed reverberating super bursts (RSB). RSBs are network events classified as single network bursts using standard burst detection algorithms. However, using our algorithm we show that RSBs are truly composed of an initial prolonged large amplitude burst followed by several mini-bursts occurring at a high frequency. This pattern resembles reverberations such as those observed in ‘epileptic’ networks^38–40^. When RSBs and the containing mini-bursts were factored in, *MECP2* null networks demonstrated increased total number of network bursts, consistent with the hyperexcitability phenotype seen in single neurons. Thus, hyperexcitable *MECP2* null neurons led to a hyperexcitability phenotype in neuronal networks.

Some non-reverberating mutant networks showed no significant differences in network burst duration relative to non-reverberating isogenic control networks. The latter suggests that there is a variability in the hyperexcitability phenotype perhaps caused by compensatory mechanisms that can restore the network phenotype even in the absence of *MECP2*. Our results highlight the importance of careful consideration when utilizing high-throughput MEA in network phenotyping. The fact that some isogenic control networks also show RSBs suggests that this is not a phenomenon only linked to *MECP2* mutant networks but may be more ubiquitous in MEA studies than previously reported^41–46^. It may be that RSBs are a normal part of the functional trajectory in a developing neuronal network that is presenting itself earlier, more often, and is unable to transition past in *MECP2* null networks.

### Mechanisms of RSBs generation

The mechanisms of rapid fluctuations in neuronal activity producing reverberations could be grouped into: 1) abnormalities in the excitation/inhibition balance^26^, 2) abnormalities in synaptic calcium dynamics^27–29,47^, and 3) an interplay between the two^25^. The former mechanism has been proposed by a study in brain organoids containing a mix of excitatory and inhibitory neurons^26^. The use of bicuculline, a GABA_A_ receptor antagonist, decreased the number of RSBs (referred to as nested oscillations in the study^26^) while preserving network bursts. Therefore, the authors proposed that the interplay between inhibitory and excitatory synapses was the likely source of RSBs. However, in our study, the networks did not contain inhibitory interneurons, and we still observed RSBs, that were more frequent in *MECP2* null neurons.

Moreover, we used bicuculline and found no effect on RSBs indicating that first, our networks were indeed excitatory-only, and second that our RSBs were not caused by the interplay between excitatory and inhibitory neurotransmission. There were subtle increases in network event frequency, possibly due to the blockage of Ca^2+^-activated potassium channels by bicuculline. This can result in increased firing frequency via blockade of the slow after-hyperpolarization phase of action potentials, and therefore faster burst frequency^35,36,48^.

On the other hand, the application of EGTA-AM had robust effects in eliminating RSBs, strongly implicating slow-kinetic calcium dynamics in maintaining RSBs. EGTA-AM can enter the synaptic terminal causing a decrease in asynchronous neurotransmitter release without affecting the large, localized Ca^2+^ transients responsible for synchronous release^37,49^. Importantly, EGTA-AM was further shown to restore burst phenotype metrics in reverberating networks to similar levels as isogenic controls. The latter may open a possibility for rescuing network hyperexcitability phenotypes during the early stages of brain development. This, however, needs further investigation due to the important role of Ca^2+^ in neuronal excitability during development.

One possible explanation for these results is that RSBs are maintained by elevated Ca^2+^ in the presynaptic terminal, which triggers asynchronous neurotransmitter release^50^. Asynchronous release occurs when Ca^2+^ levels remain elevated even after the initial stimulus has ended, typically after moderate to high-frequency stimulation^29,50^. Lau and Bi suggest that asynchronous neurotransmitter release plays crucial roles in the sustained persistence of activity after transient inputs – implicating its function in working memory and motor planning circuits and demonstrating numerous mechanisms further implicated in maintaining reverberating-like activity^25,51^. Dao Duc et al. suggested reverberating-like activity would be detectable as changes in amplitude of excitatory postsynaptic currents evoked by paired stimuli with a short inter-stimulus-interval, reminiscent of Lau and Bi’s paired-pulse protocol that replicated reverberating-like activity^25,29^. The decrease in rheobase observed in the *MECP2* null neurons of our study may also contribute to the recruitment of voltage-sensitive Ca^2+^ channels in the presynaptic terminal and its availability to trigger asynchronous neurotransmitter release^52^.

Reverberating networks had a more prolonged initiation burst duration compared to non-reverberating network mean burst duration. EGTA-AM treatment reduced initiation burst duration in reverberating networks, suggesting the association that the longer initiation burst duration may produce elevated presynaptic Ca^2+^ build-up, which would trigger reverberating mini-bursts in genetically predisposed hyperexcitable neurons. It is likely that *MECP2* null networks exist in a hyperexcitable state, such that physiologically relevant stimuli are more likely to trigger a network reverberation. RSBs may serve as a predisposing factor for the development of disorders of hyperexcitability such as epilepsy which has been shown to have numerous etiologies^39,47,53^. Reverberations can induce long-term potentiation in synapses and trigger mechanisms of synaptic plasticity^50,51,54,55^. During the early stages of network development, such mechanisms are critical for forming connections between neurons and establishing different modes of network dynamics. One possibility is that RSBs serve as a compensatory mechanism for *MECP2* null networks to achieve similar levels of connectivity as typically developed networks. However, this persistent excitation coupled with hyperexcitability may contribute to the development of seizures – a comorbidity observed in RTT patients^39,47^.

The ability of excitatory networks to generate RSBs associated with their connectivity level has also been linked to asynchronous neurotransmitter release^28,29^. As bursting patterns similar to RSBs have been previously reported in healthy, developing neuronal networks, it may be that RSBs are a normal part of network development as neurons functionally integrate more strongly with each other^21^. At early time points networks are weakly connected, and excitatory inputs may be insufficient to produce successive rounds of reverberating mini-bursts. As synaptic strength increases, this will permit noisy asynchronous neurotransmitter release to elicit RSBs. Over time, the rapid and strong initiation burst in these RSBs would deplete neurons of neurotransmitter resources, precluding additional rounds of reverberating mini-bursts and establishing mature bursting dynamics. Therefore, hypoconnected *MECP2* null networks may persist in a bursting regime that is dominated by RSBs, unable to transition into a mature network. In contrast, hyperconnected networks in other neurodevelopmental disorders may resolve RSBs more quickly than controls to establish mature networks, and may require more frequent MEA recordings for RSB detection.

Interestingly, the CLT phenotype did not show the increase in single neuron excitability nor the clear increase in network excitability^7^ observed in the *MECP2* null phenotypes (WIBR3 and PGPC14), suggesting that RSBs dominating network dynamics may be linked to the more severe *MECP2* mutations^56^. The CLT phenotype was not hyperexcitable nor showed significant differences in AMPAR-mEPSC amplitude or frequency, alongside no changes in synaptic density, whereas the *MECP2* null phenotype was reduced across the board^7^. It may be that the CLT phenotype does not have the same magnitude of changes underlying connectivity to demonstrate the differences in RSBs compared to the *MECP2* null mutants. The rescue of the hyperexcitable phenotype by EGTA-AM suggests Ca^2+^ dynamics are critical for the emergence of RSBs. Our findings demonstrate that *MECP2* null neurons exhibit hyperexcitability that can be rescued pharmacologically. If the RSB mechanism perpetuates into childhood, it may predispose RTT patients with *MECP2* null mutations to develop seizures, and provide insight into possible Ca^2+^ mediated pharmacological interventions of epilepsy.

## MATERIALS AND METHODS

### Cell culture and differentiation

Three human-derived stem cell lines were previously generated. Briefly, CLT L124W mutant iPSC lines were derived from fibroblasts of a clinically diagnosed Rett Syndrome patient. WIBR3 was derived from an unaffected ESC line, and PGPC14 was from an unaffected iPSC line. Using XCI assays, isogenic control pair for the CLT line was identified. Gene editing was used to generate isogenic null mutant pairs for the unaffected lines (WIBR3, PGPC14). *Neurogenin-2* (*Ngn2*) overexpression protocol rapidly differentiated hSC lines towards excitatory cortical neurons. All cell lines were examined for MECP2 protein levels. For further information, refer to Mok et al. (2022)^7^.

### MEA plating and recording

MEA plating was performed as previously described in Mok et al. (2022)7. Briefly, 12-Well Cytoview plates (Axion Biosystems) containing 64 electrodes per well were coated with sterilized 0.1% PEI solution in borate buffer solution pH 8.4 for 1 hour at room temperature. Plates were then washed 4 times with sterile water and allowed to dry overnight. Day 8 Ngn2 neurons were then seeded at a density of 100,000 cells per well in 100 µl droplets of CM2 Brainphys media [Brainphys (STEMCELL technologies), 1x Glutamax, 1x pen/strep, 10 ng/ml BDNF, 10 ng/ml GDNF], supplemented with 400 μg/ml laminin and 10 μM ROCK inhibitor. Seeded cells were allowed to settle for 2 hours to ensure good adherence to the recording surface before wells were carefully flooded with an additional 1 ml of CM2 Brainphys media, supplemented with 40μg/ml laminin. The following day, P1 mouse astrocytes were added to each well at a seeding density of 20,000 cells per well, maintaining a 5:1 ratio of neurons to astrocytes. Cell culture media was changed twice weekly during the recording schedule, with media changes occurring exactly 24 hours prior to each recording, with the exception of pharmacological treatment experiments (see Pharmacological treatments). For MEA recordings, each plate was allowed to incubate for 5 minutes on an Axion Maestro device heated to 37°C under 5% CO_2_. Spontaneous activity was then recorded at a sampling frequency of 12.5 kHz for 5 minutes using AxIS v2.0 software, with the analog filter settings set to “Neural: Spikes” mode (1200X gain, 200 – 5000 Hz bandwidth, median referencing). Neural spikes recordings were further bandpass filtered at 0.2 – 3 kHz and spikes were detected using a threshold crossing method with the threshold set at 6x the standard deviation of the noise of recording electrodes. Further analyses of recorded spikes were performed in Python (https://www.python.org).

### Pharmacological treatments

MEA plates were recorded twice weekly as described in MEA Plating and Recording and were monitored for the appearance of RSBs. Pharmacological treatments began after RSBs were detected in cultures. Before adding any pharmacological compounds, culture media was first changed with fresh CM2 media and plates were allowed to incubate for 1 hour to allow neuronal activity to stabilize. Baseline spontaneous activity was then recorded for 10 minutes at 37 °C. Immediately following baseline recording, one pharmacological agent (0.1% DMSO, or 10 µM bicuculline) was added to each well and 10 minutes of post-treatment spontaneous activity was recorded. Culture media was then changed 3 times to washout pharmacological agents and plates were allowed to incubate for another hour to allow activity to stabilize. A new 10-minute baseline recording was then taken, and the process was repeated until all pharmacological agents had been tested. 25 µM EGTA-AM was always added as the final drug compound in the rounds of testing as it was observed that normal baseline spontaneous activity did not return even after drug washout.

### Intracellular Adaptation

To quantify adaptation in the intracellular current clamp recordings, spike times were first detected using Allen Institute’s Intrinsic Physiology Feature Extractor (IPFX) Python package (https://github.com/AllenInstitute/ipfx). Action potential spike times across each current injection sweep was organized into 0.05s bins, producing the distribution of spike times for each neuron at all stimulation currents. Using SciPy’s non-linear least squares curve_fit() function (https://docs.scipy.org/doc/scipy/reference/generated/scipy.optimize.curve_fit.html), a mono-exponential decay function was fitted using the equation below:

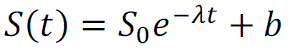

where *S(t)* represents the number of spikes at a given time (*t)*. *S*_0_ is the initial amount at *t* = 0. *e* is Euler’s constant. λ is the rate of decay. *b* represents the baseline of spiking activity, at any given time.

### ISI Threshold Burst Detection

All ISI-based burst detection was performed using Axion Biosystem’s Neural Metrics tool (v2.2.4) using AxIS (2.0) software generated AxIS Spike (.spk) files. 15ms fixed ISI burst detection used a 15ms max ISI threshold with requirements of a minimum of 50 spikes and 35% of electrodes active. 100ms fixed ISI burst detection used a 100ms max ISI threshold with requirements of a minimum of 50 spikes and 35% of electrodes active. Adaptive ISI required a minimum of 50 spikes and 35% of electrodes active. The resulting Neural Metrics feature statistics were exported to a CSV file, compiled, and then analyzed using custom-built Python code.

### Power Spectral Density Estimation

Spike times from each electrode were converted into a binary spike matrix with a bin size of 0.00008 seconds (i.e., 1/12.5kHz sampling frequency). Each electrode’s spike matrix then underwent power spectral density (PSD) estimate using SciPy’s welch() function (https://docs.scipy.org/doc/scipy/reference/generated/scipy.signal.welch.html) and expressed in decibels (dB) as a ratio against the PSD of a randomly permuted version of the same spike matrix. Each channel’s PSD was averaged to generate the network-averaged PSD, representing the network’s frequency components. Peaks in the spectra between 0Hz and 1Hz typically represented network burst frequency, whereas mini-burst frequency was expected to be between 1Hz and 10Hz.

### Spike sorting

Individual neuronal units were identified from extracellular multielectrode arrays according to Axion Biosystem’s Spike Sorting Protocol^31,57^. In brief, multi-unit action potentials were extracted from raw voltage signals using AxIS Navigator (Axion Biosystems) and stored as an AxIS Spike (.spk) file (see MEA Plating and Recording). AxIS Spike files were converted to NeuroExplorer (.nex) file using Axion Data Export Tool. NeuroExplorer files were loaded into Plexon Offline Sorter (version 4.5.0). All channels underwent Automatic Sorting using a K-Means Scan with a range of 1 to 6. All spike sorting results underwent manual quality control^31^.

### Reverberating Super Burst Detection

#### See Supplementary Figure 5 for the algorithm block diagram. Signal Conditioning

Spike times from each channel were converted to a binary spike matrix with bin sizes of 0.00008 seconds (i.e., 1/12.5kHz sampling frequency). Each spike matrix was convolved with a bandwidth-optimized Gaussian kernel^30^ to generate a spike density function, representing the estimated instantaneous firing rate of each channel. The average instantaneous firing rate of the network was calculated by computing the spike count weighted average spike density function across all electrodes. Networks that did not exhibit a maximum instantaneous firing rate greater than 3Hz were excluded from the analysis, presumed to be non-bursting wells.

Network bursts were identified as having local maxima with a peak amplitude that exceeded a minimum burst threshold (10% of the maximum firing rate with a minimum of 3Hz). Additionally, burst peaks required a prominence of 50% of the maximum firing rate of the network. If the number of burst peaks exceeded 5 bursts, the analysis would proceed to the *Burst Detection Loop* and *Reverberating Network Detection*.

#### Burst Detection Loop

Preliminary burst boundaries, defining the start and end of network bursts, were calculated by finding the local maxima and minima of the first derivative of the spike density function. Local maxima represented the start of the burst, whereas local minima represented the end of the burst. Burst boundaries were assigned to previously calculated burst peaks that fell between the boundaries.

#### Reverberating Network Detection

To identify reverberating networks, K-means clustering (and later Gaussian Mixed Models) was performed with amplitude (i.e., maximum firing rate) in *n*-1 burst and inter-burst-peak-interval between each burst (*n*).

Reverberating networks had two clearly defined clusters. The first cluster localized to the high inter-burst-peak-interval and high firing rate amplitude dimensions – representing the initiation network burst. The second cluster localized to the low inter-burst-peak interval and low firing rate amplitude dimensions – representing the mini-bursts.

Non-reverberating networks would exhibit one of the following:

1. One cluster: demonstrating a network that was highly periodic with low variability in the maximum amplitude of firing rate in each burst.
2. Two clusters: demonstrating a noisy network that contained maximum amplitude of firing rates that were highly variable, but often had periodic inter-burst-peak interval.

Based on this, the determining factor of a noisy versus reverberating network was minimal overlap across the both maximum firing rate and inter-burst-peak-interval dimensions, which was set to 20% overlap tolerance. Reverberating networks then proceeded to the *Reverberating Super Burst Construction Loop*.

#### Reverberating Super Burst Construction Loop

R_max_, representing the maximum inter-burst-peak-interval within RSBs, was calculated by determining the local minima between the bimodal peaks seen along the inter-burst-peak-interval distribution of a reverberating network. A_max_, representing the maximum firing rate of mini-bursts within RSBs, was calculated by determining the local minima between the bimodal peaks seen along the peak burst firing rate dimension.

Using the preliminary burst boundaries for all bursts that was previous calculated, and R_max_ and A_max_ parameters, new burst boundaries for RSBs were determined as followed. The start of every potential RSB always began with a network burst that had an amplitude exceeding A_max_. Once the initiation burst was found, if the difference between the start of the subsequent burst and the end of the current burst (i.e., inter-burst-peak-interval) was less than R_max_, the subsequent burst was included into the RSB. This was repeated until the inter-burst-peak-interval was greater than R_max_ and the end of the RSB was marked.

### RSB Feature Analysis

The following are brief explanations for select features:

#### Initiation/First Burst Duration

Calculation of the first burst duration was non-trivial due to the variability in the firing rate pattern of the initiation network burst and the proceeding reverberating mini-bursts. For example, some initiation network bursts do not reach an arbitrarily low burst threshold until the end of the RSB, whereas others do, making traditional approaches to identifying boundaries for bursts inadequate. To address this, we used the first derivative of the spike density function and identified local peaks and troughs which corresponded to changes in the firing rate velocity of the bursts, specifically the start and end of bursts, respectively. First burst duration was calculated by taking the time between the start and end of the initiation network burst.

#### Network Reverberation Index

Network Reverberation Index (NRI) was calculated for each reverberating developmental week using the following formula:

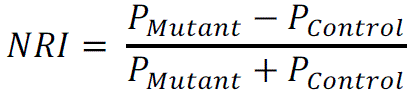

where *P* represents the proportion of reverberating networks for either mutant or control groups per individual per developmental week. NRI ∈ (–1, 1) and is positive when the mutant group is reverberating more and negative when the control group is reverberating more.

#### Network Event/Burst Duration

A network event was described as the entire RSBs (initiation network burst and mini-bursts) in reverberating networks or discrete single network bursts in non-reverberating networks. Network event duration was calculated as the average difference in time between the beginning and end of the boundaries of the network event for each well’s recording.

#### Network Event/Burst Frequency

Network event frequency was calculated as the inverse of average inter-network-event-interval between subsequent network events.

#### Number of mini-bursts/RSB

The number of mini-bursts was calculated by counting non-initiation network burst peaks within the boundaries of the RSB.

#### RSB duration

Similar to *Network Event Duration*, RSB duration was calculated by calculating the difference in time between the beginning and end of the boundaries of each RSB.

#### Mini-burst frequency

Mini-burst frequency was calculated as the inverse of inter-mini-burst-interval.

### RANSAC (RANdom Sample Conensus) Regression

Ordinary Least Squares (OLS) regression is sensitive to outliers. To generate a robust linear model for data that may contain outliers, Scikit-Learn’s RANSACRegressor() function was used (https://scikit-learn.org/stable/auto_examples/linear_model/plot_ransac.html). RANSAC sampled a subset of the data (*min_samples = 50*), fitted a linear model (*estimator = LinearRegression()*) to the selected subset of data, evaluated the model (based on the number of inliers) and then repeated for a fixed number of iterations (*max_trials=100*). The model with the best consensus score (i.e., the number of points that fall within a threshold range and are therefore considered inliers) was selected. Where used, outlier data were plotted with increased transparency and excluded in the fit. Inlier data was plotted with decreased transparency and used to generate a linear fit of the data.

### Statistical Analyses

Statistical tests for all analyses were performed on Python (3.9 and 3.11) and JASP (0.17.1). Comparisons between groups were conducted by One-way ANOVA, Two-way repeated measures ANOVA, Kruskal-Wallis, Mann-Whitney, Wilcoxon Matched Paired Signed Rank, Independent Samples Student T, and Paired Student T tests, with associated parametric and non-parametric post hoc tests where appropriate (\**p* < 0.05, \*\**p* < 0.01, \*\*\**p* < 0.001). Mono-exponential decay fit, Pearson Correlation, Linear regression, and RANSAC regression were used to identify best fits and correlations. All figures were created using Seaborn (https://seaborn.pydata.org), and Matplotlib (https://matplotlib.org). Schematics created with <u>BioRender.com</u>.

### Data Availability (NDD-Ephys-dB)

The data can be found on NDD-Ephys-dB. The NDD-Ephys-dB is a resource for electrophysiological datasets and recordings explicitly related to human neurodevelopmental disorders, including Rett Syndrome (RTT). The database contains in vitro data, including intracellular (.abf or .nwb file format) and extracellular (in Axion .spk, .csv, and HDF5 file formats) electrophysiological recordings. In addition to serving as a repository for electrophysiological data, the database provides tools like burst detection algorithms, visualization GUIs, and feature calculation toolboxes to assist with the exploration and investigation of data. NDD-Ephys-dB aims to promote data-sharing of published results and data-mining by computational neuroscientists. The goal is to expand the resource to include 3D and high-density MEA results to advance studies of all neurodevelopmental disorders.

## ACKNOWLEDGEMENTS

This research was funded by grants from SFARI (Research grant #514918 to J.E. and J.M.-T.), CIHR ERARE Team Grant (ERT161303 to J.E.), CIHR Project grant (PJT168905 to J.E.), CIHR Foundation grant (FDN-154336 to M.W.S), John Evans Leadership Fund & Ontario Research Fund (to J.E.), Canada Research Chair in Stem Cell Models of Childhood Disease (to J.E.), Beta Sigma Phi International Endowment Fund (to J.E.), BrainsCAN at Western University through the Canada First Research Excellence Fund (CFREF) (to K.S.P., JM.-T). Trainee support was provided by NSERC Postgraduate Scholarship–Doctoral (PGS-D) Scholarship (to K.S.P.), Autism Speaks Predoctoral Award and David Stephen Cant Scholarship in Stem Cell Research (to F.P.M.). We thank Rebecca Mok for providing data to NDD-Ephys-dB.

## AUTHOR CONTRIBUTIONS

Conceptualization, K.S.P., F.P.M., M.K., J.E., J.M.T.; Methodology, K.S.P., F.P.M., W.W., J.E., J.M.T.; Formal Analysis, K.S.P.; Investigation, F.P.M., W.W., W.Z.; Resources, F.P.M., W.W.; Data Curation, K.S.P., F.P.M., M.K.; Writing – Original Draft, K.S.P., F.P.M., M.K., J.E., J.M.T.; Writing – Review & Editing, K.S.P., F.P.M., M.K., W.Z., M.W.S., J.E., J.M.T.; Visualization, K.S.P.; Supervision, J.E., J.M.T.; Funding Acquisition, K.S.P., F.P.M., M.W.S., J.E., J.M.T. All authors revised and approved the paper.

## SUPPLEMENTARY FIGURES

**Supplementary Figure 1.**
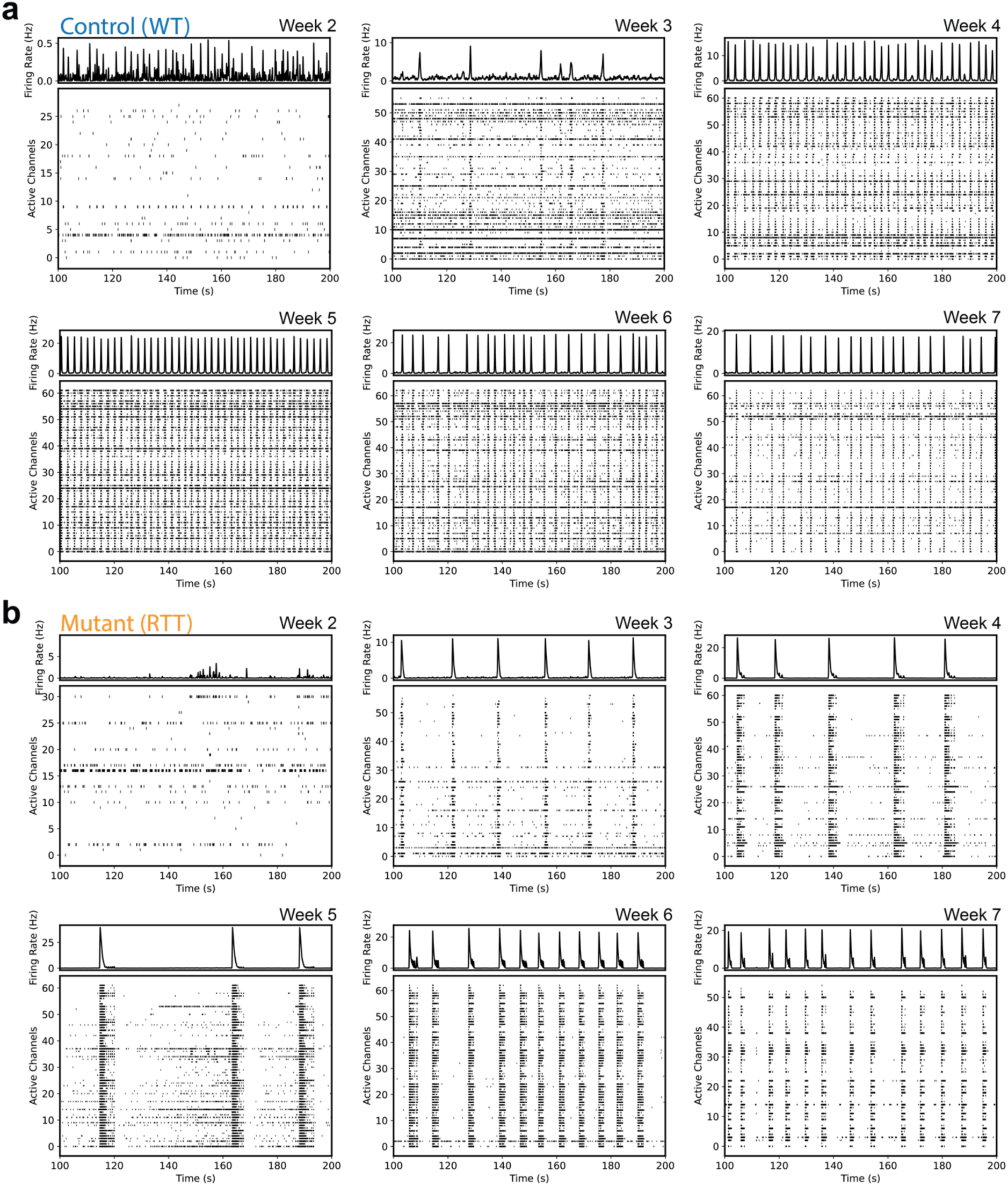
Development of *MECP2* WT and mutant networks. (a-b) Raster plots for each developmental time point from representative (a) wildtype and (b) *MECP2* mutant networks. The top subplot is the network-averaged spike density function representing the estimated instantaneous firing rate of the network.

**Supplementary Figure 2.**
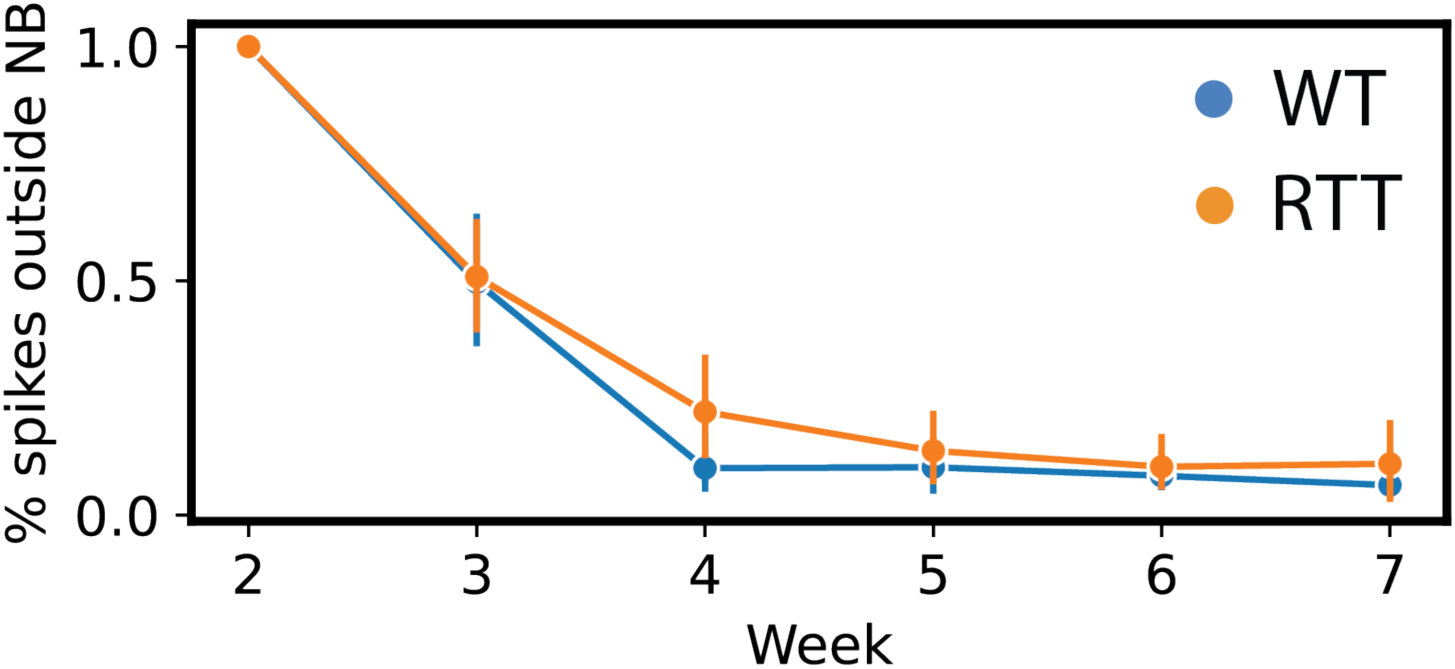
The proportion of inter-burst spiking activity. *MECP2* mutant and wildtype networks. Network activity first formed with a high degree of inter-burst spiking activity that decreased over development.

**Supplementary Figure 3.**
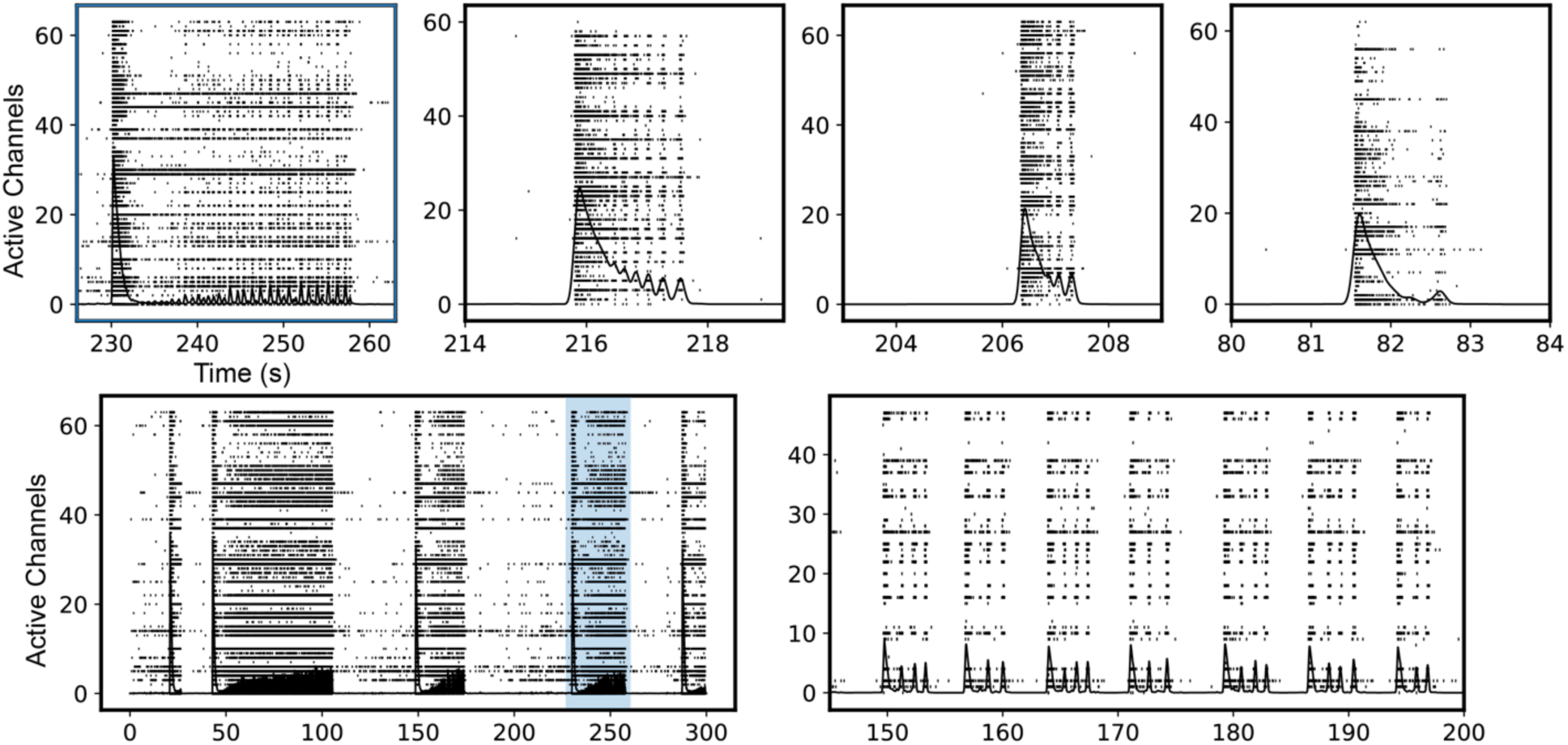
Selected examples of spike density function overlaid on raster plots demonstrating the diversity of reverberating super burst patterns.

**Supplementary Figure 4.**
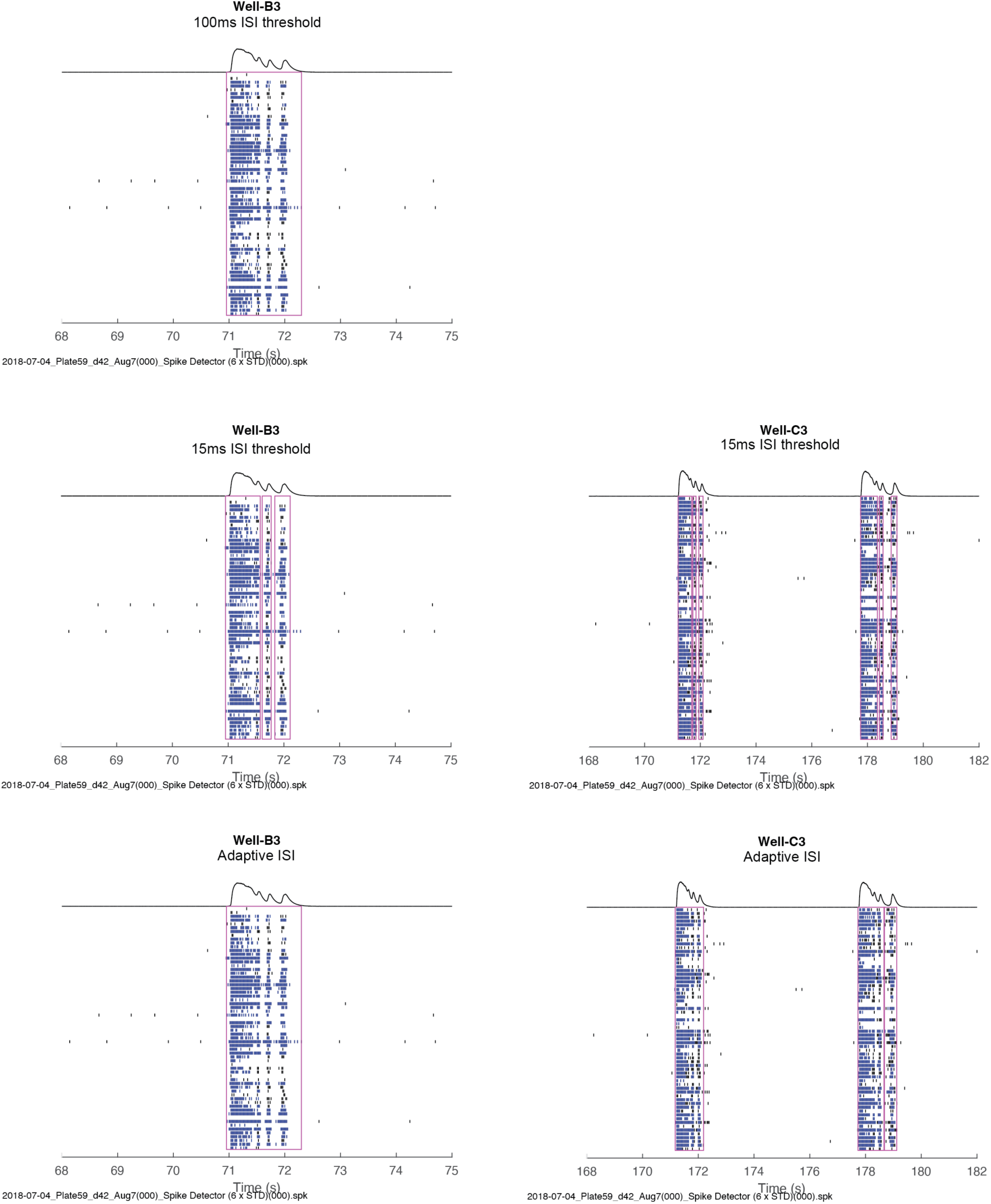
Inconsistent burst boundaries by ISI-based burst detection algorithms.

**Supplementary Figure 5.**
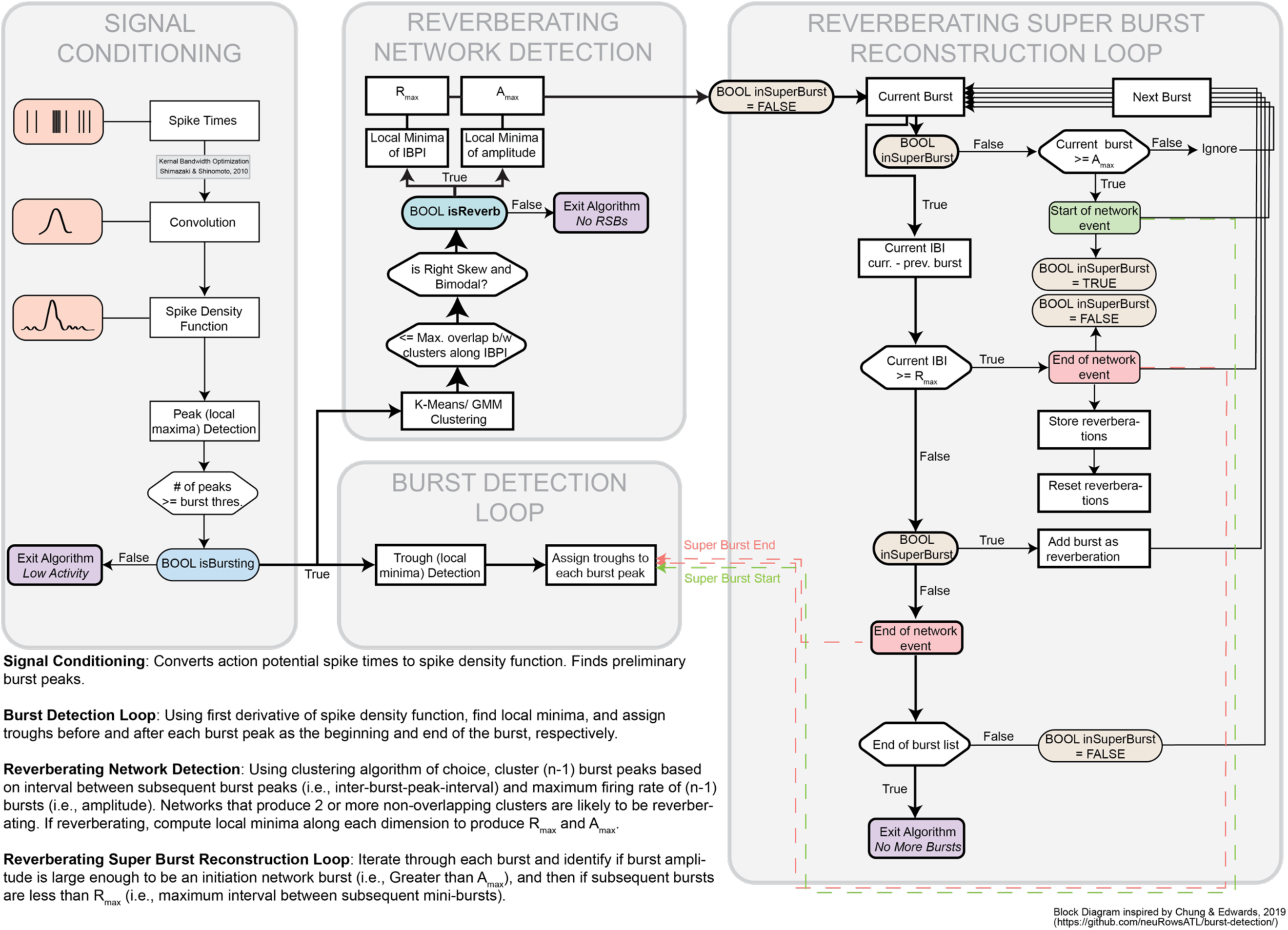
Block diagram of Burst Reverberation Detection algorithm.

**Supplementary Figure 6.**
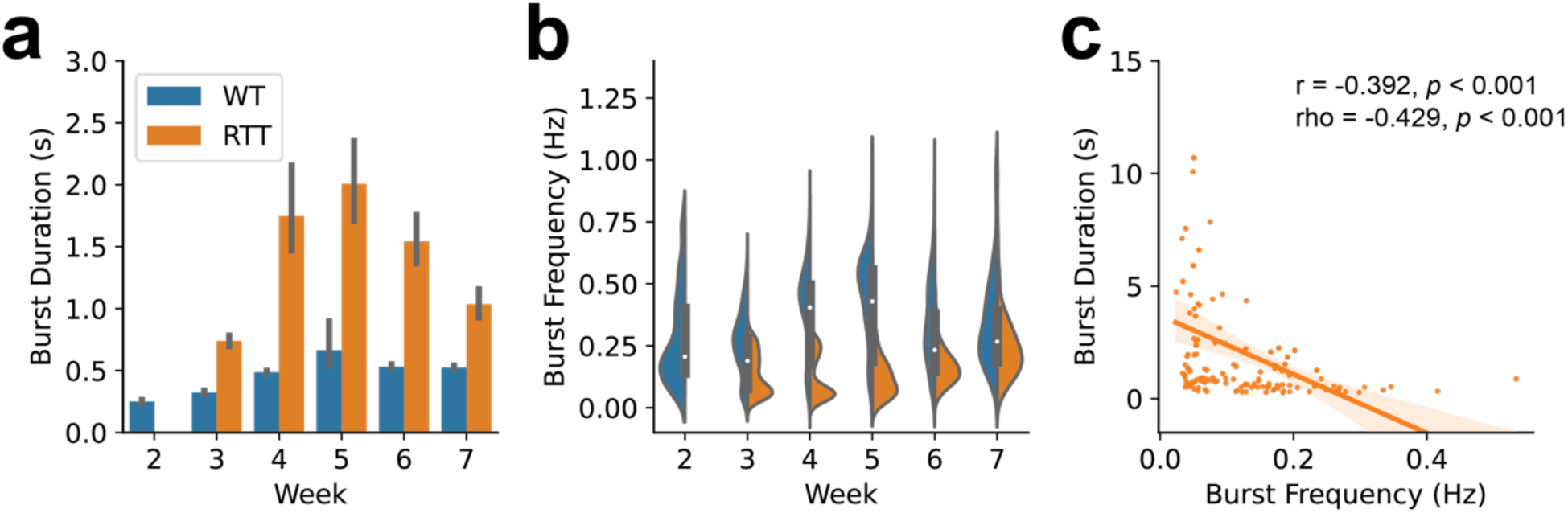
Replication of Mok et al. (2022) with SDF-based Network Burst Detection.

## REFERENCES

1. Banerjee, A., Castro, J. & Sur, M. Rett Syndrome: Genes, Synapses, Circuits, and Therapeutics. Front. Psychiatry 3, (2012).

2. Amir, R. E. et al. Rett syndrome is caused by mutations in X-linked MECP2, encoding methyl-CpG-binding protein 2. Nat. Genet. 23, 185–188 (1999).

3. Ip, J. P. K., Mellios, N. & Sur, M. Rett syndrome: insights into genetic, molecular and circuit mechanisms. Nat. Rev. Neurosci. 19, 368–382 (2018).

4. Vashi, N. & Justice, M. J. Treating Rett syndrome: from mouse models to human therapies. Mamm. Genome 30, 90–110 (2019).

5. Cardoza, B. et al. Epilepsy in Rett syndrome: Association between phenotype and genotype, and implications for practice. Seizure 20, 646–649 (2011).

6. Operto, F. F., Mazza, R., Pastorino, G. M. G., Verrotti, A. & Coppola, G. Epilepsy and genetic in Rett syndrome: A review. Brain Behav. 9, e01250 (2019).

7. Mok, R. S. F. et al. Wide spectrum of neuronal and network phenotypes in human stem cell-derived excitatory neurons with Rett syndrome-associated MECP2 mutations. Transl. Psychiatry 12, 1–16 (2022).

8. McCready, F. P., Gordillo-Sampedro, S., Pradeepan, K., Martinez-Trujillo, J. & Ellis, J. Multielectrode Arrays for Functional Phenotyping of Neurons from Induced Pluripotent Stem Cell Models of Neurodevelopmental Disorders. Biology 11, 316 (2022).

9. Deneault, E. et al. CNTN5-/+or EHMT2-/+human iPSC-derived neurons from individuals with autism develop hyperactive neuronal networks. eLife 8, e40092.

10. Nageshappa, S. et al. Altered neuronal network and rescue in a human MECP2 duplication model. Mol. Psychiatry 21, 178–188 (2016).

11. Cabrera-Garcia, D. et al. Early prediction of developing spontaneous activity in cultured neuronal networks. Sci. Rep. 11, 20407 (2021).

12. Kamioka, H., Maeda, E., Jimbo, Y., Robinson, H. P. C. & Kawana, A. Spontaneous periodic synchronized bursting during formation of mature patterns of connections in cortical cultures. Neurosci. Lett. 206, 109–112 (1996).

13. Opitz, T., De Lima, A. D. & Voigt, T. Spontaneous Development of Synchronous Oscillatory Activity During Maturation of Cortical Networks In Vitro. J. Neurophysiol. 88, 2196–2206 (2002).

14. Tozzi, A., Zare, M. & Benasich, A. A. New Perspectives on Spontaneous Brain Activity: Dynamic Networks and Energy Matter. Front. Hum. Neurosci. 10, (2016).

15. Ruthazer, E. S., Akerman, C. J. & Cline, H. T. Control of axon branch dynamics by correlated activity in vivo. Science 301, 66–70 (2003).

16. Kerschensteiner, D. Spontaneous network activity and synaptic development. Neurosci. Rev. J. Bringing Neurobiol. Neurol. Psychiatry 20, 272–290 (2014).

17. Luhmann, H. J., et al. Spontaneous Neuronal Activity in Developing Neocortical Networks: From Single Cells to Large-Scale Interactions. Front. Neural Circuits 10, (2016).

18. Zeldenrust, F., Wadman, W. J. & Englitz, B. Neural Coding With Bursts—Current State and Future Perspectives. Front. Comput. Neurosci. 12, (2018).

19. Izhikevich, E. M. Neural excitability, spiking and bursting. *Int. J*. Bifurc. Chaos 10, 1171– 1266 (2000).

20. Belykh, I., de Lange, E. & Hasler, M. Synchronization of Bursting Neurons: What Matters in the Network Topology. Phys. Rev. Lett. 94, 188101 (2005).

21. Wagenaar, D. A., Pine, J. & Potter, S. M. An extremely rich repertoire of bursting patterns during the development of cortical cultures. BMC Neurosci. 7, 11 (2006).

22. Boggio, E. M., Lonetti, G., Pizzorusso, T. & Giustetto, M. Synaptic Determinants of Rett Syndrome. Front. Synaptic Neurosci. 2, 28 (2010).

23. Della Sala, G. & Pizzorusso, T. Synaptic plasticity and signaling in rett syndrome. Dev. Neurobiol. 74, 178–196 (2014).

24. Fukuda, T., Itoh, M., Ichikawa, T., Washiyama, K. & Goto, Y. Delayed maturation of neuronal architecture and synaptogenesis in cerebral cortex of Mecp2-deficient mice. J. Neuropathol. Exp. Neurol. 64, 537–544 (2005).

25. Lau, P.-M. & Bi, G.-Q. Synaptic mechanisms of persistent reverberatory activity in neuronal networks. Proc. Natl. Acad. Sci. U. S. A. 102, 10333–10338 (2005).

26. Trujillo, C. A. et al. Complex Oscillatory Waves Emerging from Cortical Organoids Model Early Human Brain Network Development. Cell Stem Cell 25, 558–569.e7 (2019).

27. Huang, C.-H., Huang, Y.-T., Chen, C.-C. & Chan, C. K. Propagation and synchronization of reverberatory bursts in developing cultured networks. J. Comput. Neurosci. 42, 177–185 (2017).

28. Volman, V., Gerkin, R. C., Lau, P.-M., Ben-Jacob, E. & Bi, G.-Q. Calcium and synaptic dynamics underlying reverberatory activity in neuronal networks. Phys. Biol. 4, 91–103 (2007).

29. Dao Duc, K., et al. Bursting Reverberation as a Multiscale Neuronal Network Process Driven by Synaptic Depression-Facilitation. PLOS ONE 10, e0124694 (2015).

30. Shimazaki, H. & Shinomoto, S. Kernel bandwidth optimization in spike rate estimation. J. Comput. Neurosci. 29, 171–182 (2010).

31. Corrigan, B. W. et al. Distinct neural codes in primate hippocampus and lateral prefrontal cortex during associative learning in virtual environments. Neuron 110, 2155–2169.e4 (2022).

32. Cotterill, E., Charlesworth, P., Thomas, C. W., Paulsen, O. & Eglen, S. J. A comparison of computational methods for detecting bursts in neuronal spike trains and their application to human stem cell-derived neuronal networks. J. Neurophysiol. 116, 306–321 (2016).

33. Pasquale, V., Martinoia, S. & Chiappalone, M. A self-adapting approach for the detection of bursts and network bursts in neuronal cultures. J. Comput. Neurosci. 29, 213–229 (2010).

34. Bakkum, D. J. et al. Parameters for burst detection. Front. Comput. Neurosci. 7, (2014).

35. Khawaled, R., Bruening-Wright, A., Adelman, J. P. & Maylie, J. Bicuculline block of small-conductance calcium-activated potassium channels. Pflugers Arch. 438, 314–321 (1999).

36. Johansson, S., Druzin, M., Haage, D. & Wang, M. D. The functional role of a bicuculline-sensitive Ca2+-activated K+ current in rat medial preoptic neurons. J. Physiol. 532, 625–635 (2001).

37. Nakamura, Y. EGTA Can Inhibit Vesicular Release in the Nanodomain of Single Ca2+ Channels. Front. Synaptic Neurosci. 11, 26 (2019).

38. Blumenfeld, H. Cellular and Network Mechanisms of Spike-Wave Seizures. Epilepsia 46, 21–33 (2005).

39. McCormick, D. A. Persistent Cortical Activity: Mechanisms of Generation and Effects on Neuronal Excitability. Cereb. Cortex 13, 1219–1231 (2003).

40. Sanchez-Vives, M. V. & McCormick, D. A. Cellular and network mechanisms of rhythmic recurrent activity in neocortex. Nat. Neurosci. 3, 1027–1034 (2000).

41. Droge, M., Gross, G., Hightower, M. & Czisny, L. Multielectrode analysis of coordinated, multisite, rhythmic bursting in cultured CNS monolayer networks. J. Neurosci. 6, 1583–1592 (1986).

42. Pimashkin, A. et al. Spiking Signatures of Spontaneous Activity Bursts in Hippocampal Cultures. Front. Comput. Neurosci. 5, (2011).

43. Heikkilä, T. J. et al. Human embryonic stem cell-derived neuronal cells form spontaneously active neuronal networks in vitro. Exp. Neurol. 218, 109–116 (2009).

44. Matsuda, N. et al. Detection of synchronized burst firing in cultured human induced pluripotent stem cell-derived neurons using a 4-step method. Biochem. Biophys. Res. Commun. 497, 612–618 (2018).

45. Odawara, A., Katoh, H., Matsuda, N. & Suzuki, I. Physiological maturation and drug responses of human induced pluripotent stem cell-derived cortical neuronal networks in long-term culture. Sci. Rep. 6, 26181 (2016).

46. Iida, S., Shimba, K., Sakai, K., Kotani, K. & Jimbo, Y. Synchronous firing patterns of induced pluripotent stem cell-derived cortical neurons depend on the network structure consisting of excitatory and inhibitory neurons. Biochem. Biophys. Res. Commun. 501, 152– 157 (2018).

47. Holcman, D. & Tsodyks, M. The Emergence of Up and Down States in Cortical Networks. PLoS Comput. Biol. 2, (2006).

48. Rhoades, B. K. & Gross, G. W. Potassium and calcium channel dependence of bursting in cultured neuronal networks. Brain Res. 643, 310–318 (1994).

49. Adler, E., Augustine, G., Duffy, S. & Charlton, M. Alien intracellular calcium chelators attenuate neurotransmitter release at the squid giant synapse. J. Neurosci. 11, 1496–1507 (1991).

50. Kaeser, P. S. & Regehr, W. G. Molecular Mechanisms for Synchronous, Asynchronous, and Spontaneous Neurotransmitter Release. Annu. Rev. Physiol. 76, 333–363 (2014).

51. Iremonger, K. J. & Bains, J. S. Integration of Asynchronously Released Quanta Prolongs the Postsynaptic Spike Window. J. Neurosci. 27, 6684 (2007).

52. Catterall, W. A. Voltage-Gated Calcium Channels. Cold Spring Harb. Perspect. Biol. 3, a003947 (2011).

53. Staley, K. Molecular mechanisms of epilepsy. Nat. Neurosci. 18, 367–372 (2015).

54. Lu, T. & Trussell, L. O. Inhibitory Transmission Mediated by Asynchronous Transmitter Release. Neuron 26, 683–694 (2000).

55. Hjelmstad, G. O. Interactions Between Asynchronous Release and Short-Term Plasticity in the Nucleus Accumbens Slice. J. Neurophysiol. 95, 2020–2023 (2006).

56. Pintaudi, M. et al. Epilepsy in Rett syndrome: Clinical and genetic features. Epilepsy Behav. 19, 296–300 (2010).

57. Spike Sorting Protocol | Axion Biosystems. https://www.axionbiosystems.com/resources/culture-protocol/spike-sorting-protocol.

